# Natural Product-Like Fragments Unlock Novel Chemotypes for a Kinase Target – Exploring Options beyond the Flatland

**DOI:** 10.1101/2025.06.11.659015

**Authors:** Anna Santura, Janis Müller, Madita Wolter, Ina-Charlotte Tutzschky, Moritz Ruf, Alexander Metz, Anna Sandner, Stefan Merkl, Gerhard Klebe, Serghei Glinca, Paul Czodrowski

**Author notes:** A.S. is the main contributor of the manuscript. All authors have given approval to the final version.

## Abstract

In this study we utilized a high-performance soaking system of protein kinase A (PKA) to perform a crystallographic screening of a natural product-like fragment library. We resolved 36 fragment-bound structures, corresponding to a hit rate of 41%. Nine fragments bound within the ATP site, nine peripherally, and 18 interacted with both the ATP and peripheral sites. One fragment binds to the same site as the approved allosteric kinase inhibitor asciminib, while another induces an unexpected conformational change. Systematic database mining revealed that both the fragments and their natural product parents have not been previously associated with PKA or kinase activity. A scaffold/chemotype analysis further underscored their novelty. Cheminformatics analyses confirmed that these fragments occupy a distinct chemical space, enriched in saturation, spatial complexity and molecular three-dimensional character compared to kinase binders from reference datasets. These properties have previously been linked to increased selectivity, reduced CYP450 inhibition, and higher overall clinical success rates.

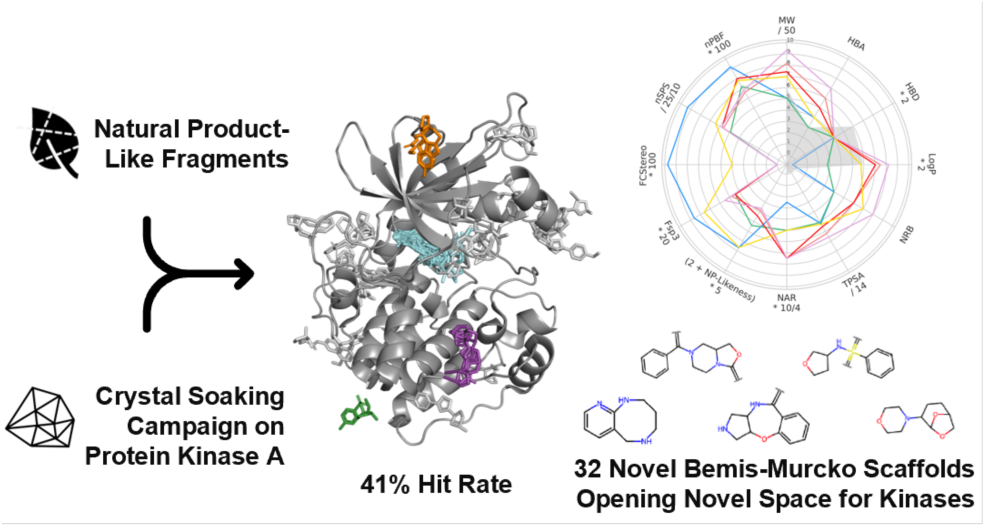

## 1 Introduction

### 1.1 Fragment Screening Libraries Benefit From Increased Molecular Three-Dimensionality

The success of fragment-based drug discovery (FBDD) heavily depends on the quality and structural variation, including the coverage of distinct chemotypes, of the fragment library.^1^ Although chemical diversity in FBDD libraries has been achieved in many respects, considering a large variety of shapes – particularly in terms of fragments rich in three-dimensional (3D) character – remains a key challenge.^2–4^ In the past, many fragment libraries exhibited a preponderance of sp2-hybridized carbons, aromatic and heterocyclic rings, as a result of synthetic methods favoring sp2-sp2 bond formation.^5^ However, the scope is changing and there is growing interest in, and a strong rationale for the inclusion of spatially more 3D building blocks in fragment libraries.^2,4^ Recent efforts have addressed synthetic challenges associated with the synthesis (and elaboration) of such sp^3^-rich fragments, and thus focused predominantly on tailored library design, ready for the screening against biological targets.^4,6^

Compound shape has long been recognized as the primary factor for successful binding of a ligand and to its biological target(s). Incorporating out-of-plane substituents and starting with spatial core structures can optimize ligand-target complementarity. The spatial orientation of pharmacophoric features is essential for effective target binding and optimizing protein-ligand interactions, thus potentially improving potency and selectivity while mitigating off-target effects. Indeed, systematic studies by Lovering *et al.*^7,8^ demonstrated that an increased fraction of sp^3^-hybridized/chiral carbons enhances selectivity and avoids cytochrome P450 inhibition, a key factor to control and design metabolism. Molecules incorporating significant 3D character often display improved aqueous solubility due to less efficient and therefore overall weaker solid-state crystal lattice packing.^9^ Taken together, an enhanced 3D character may contribute positively to both improved pharmacodynamics and pharmacokinetics, and has been correlated to overall success of drug candidates in clinical studies.^7^ At the same time, it can be argued that the increase in molecular complexity may reduce hit rates in screening campaigns.^10–12^ Whilst the cited studies do not apply to FBDD approaches specifically, it is reasonable to propose that 3D fragments could be valuable starting points for drug development.^4^

While the 2009 publication by Lovering *et al.*^7^ was expected to be a turning point in medicinal chemistry, efforts in the last years mostly addressed the synthetic challenges associated with sp^3^-rich fragments. However, from a medicinal chemistry perspective, a re-evaluation of the 15-year-old ‘*Escape from Flatland* ‘ paper did not find a clear relationship between the highest (pre-)clinical development phase reached by a compound and its fraction of sp^3^-hybridized carbons after 2009 anymore.^13^ This apparent disconnect in the last years was attributed to the changing trend in drug target selection, in particular an increase in the number of kinase inhibitors, as well as an increase in the use of metal-catalyzed cross coupling reactions simplifying the incorporation of sp2-hybridized systems into drugs.^13^ As described, however, there is still a strong rationale and promising perspective for the inclusion of sp^3^-rich fragments in screening libraries, also for kinase targets.

Given the inherent richness of natural products (NPs) in saturation, sp^3^-hybridized carbons, and stereo centers compared to synthetic molecules,^14^ NP-inspired molecules offer a promising strategy for creating shape-diverse fragment libraries.^15–18^ Additionally, NPs typically contain fewer aromatic rings which potentially enhances aqueous solubility, and suffer less from properties falling under the pan-assay interference compounds.^14,19^ In other words, NPs as well as NP-derived fragments occupy areas of the chemical space likely underexplored by molecules obtained by classical organic synthesis.^20^ A proof-of-concept study by Over *et al.*^15^ demonstrated that NP-derived fragments can give access to novel chemotypes for established drug targets, such as p38α kinase. Despite this success, NPs have so far received only little attention as a pool for fragment-based screening primarily due to their synthetic intractability and the challenging chemistry required for structural expansion and optimization.^18,21^ More recently, Huschmann *et al.*^18^ highlighted the utility of a commercially available library of NP-derived fragments for initial crystallographic screening, and subsequent hit validation using off-the-shelf follow-up compounds.

### 1.2 Kinases are in the Focus of Drug Discovery

Protein kinases are among the largest protein families in the human genome, comprising 518 members.^22^ As approximately 30% of all human protein kinases are associated with disease, kinases are in the focus of the pharmaceutical industry and academic research.^23^ Most protein kinases share a very similar 3D structure, the so-called eukaryotic protein kinase fold. Among them, protein kinase A (PKA), also known as cAMP-dependent protein kinase, stands out as one of the best-studied and well-characterized protein kinase to date.^24,25^ As PKA is a well-established system for crystallization and qualifies for the ‘crystal structure first approach’,^26,27^ it can serve as a surrogate to study other members of the protein kinase family. Notably, the structure of the PKA catalytic subunit α was the first kinase structure to be determined.^28^ As of today, more than 5000 crystal structures comprising the kinase catalytic domain have been deposited in the Protein Data Bank (PDB),^29^ thereof about 350 structures of PKA, and many more are available in proprietary databases.

As of September 2024, 107 small molecule protein kinase inhibitors (PKIs) targeting around 20 different protein kinases have been approved, primarily for treating neoplasms, and many more are currently in clinical trials worldwide.^30–32^ Most PKIs currently on the market, target the highly conserved and ubiquitously present binding site of the adenine triphosphate (ATP) cofactor, making selectivity difficult to achieve as difference between kinases are found only in the back and front pocket or above and below the ATP binding site. Most orthosteric PKIs mimic the cofactor’s adenine ring by forming 1–3 hydrogen bonds with the kinase hinge motif. This conserved interaction pattern is reflected in most chemotypes of the orthosteric inhibitors, which predominantly correspond to (flat, aromatic) *N*-heterocycles.^33^ The nonselective natural product staurosporine and its derivatives also address the hinge motif and inhibit kinases which underscores that sterically more demanding compounds are amenable to modulate kinase activity.^33,34^ Notably, the introduction of sterically more complex metal organic centers provide the promiscuous staurosporine scaffold with remarkable selectivity features. Despite such systematic studies have shown the superiority of shapely sp^3^-rich compounds, even today kinase-targeted screening libraries are dominated by flat heteroaromatic compounds. As a consequence, presently the FDA-approved kinase inhibitors all possess a minimum of two aromatic rings and exhibit an assembly of rather planar building blocks.^30,35^

PKIs binding to so-called allosteric binding sites are generally expected to offer a selectivity advantage compared to orthosteric PKIs, as they mostly bind to sites that are less conserved across the kinome. Additionally, such PKIs are thought to be less prone to the development of resistance. Recent systematic analyses of publicly available kinase ligand complex structures mapped in total 12 binding pockets in the kinase domain (designated A–L), including the ATP site (A) and 11 peripheral binding pockets (B–L), although not all are present in every kinase.^36,37^ A compelling example of a successful allosteric PKIs is asciminib, which specifically targets the E pocket of BCR-ABL1 kinase.^38^ Approved in 2021 for the treatment of chronic myelogenous leukemia, asciminib is effective against both wildtype BCR-ABL1 and several mutated forms, thus meeting all the promises for allosteric PKIs.^38^

As with drug candidates for other target classes, enhancing molecular three-dimensionality in general, and introducing carbon bond saturation and stereogenic centers in particular, may help to escape from flatland and to secure new intellectual property of future PKIs.^7,31^ Isolated examples of screening 3D-rich fragments against a kinase target have been published.^39,40^

### 1.3 Objectives

This FBDD study aimed to address the challenge of enhancing molecular three-dimensionality in general, and introducing carbon bond saturation and stereogenic centers in specific, particularly for kinase targets. By utilizing a a set of fragments from the ‘Fragments from Nature’ library, we sought to leverage the inherent spatial complexity of NP-inspired fragments, to identify promising starting points for the development of new PKIs. In detail, we conducted a crystal soaking campaign on protein kinase A (PKA) using NP-like fragments, adhering to the ‘crystal structure first’ paradigm. Our structure-based analyses focused on classifying the kinase conformation, including the conformation of the glycine-rich loop; on the fragment binding sites, with emphasis on the peripheral sites; and the discovery of potential novel kinase binding modalities. We further complemented these efforts with a systematic database mining to contextualize the identified fragment hits within the broader landscape of NPs and kinase ligands. Specifically, we assessed whether the bound fragments indeed represent novel scaffolds, and quantified the degree of spatial complexity shown by the fragments compared to reference molecules.

## 2 Results and Discussion

### 2.1 Fragment Library and Crystal Soaking Campaign

A selection of 87 fragments from AnalytiCon Discovery’s FRGx (’Fragments from Nature’) library^41^ was utilized in a crystal soaking campaign on PKA. The campaign delivered an exceptionally high hit rate of 41%, resulting in 36 high-resolution complex structures. All binding events were identified in the native electron density map and all structures could be fully refined. All complex structures were resolved at a resolution below 2 Å, with an average resolution of 1.5 Å. In 61% of the complex structures more than one fragment copy binds to the protein. Considering all of these copies, a total of 69 distinct binding poses were determined. The chemical structures of the 36 fragment hits are visualized in Fig. 1.

**Figure 1:**
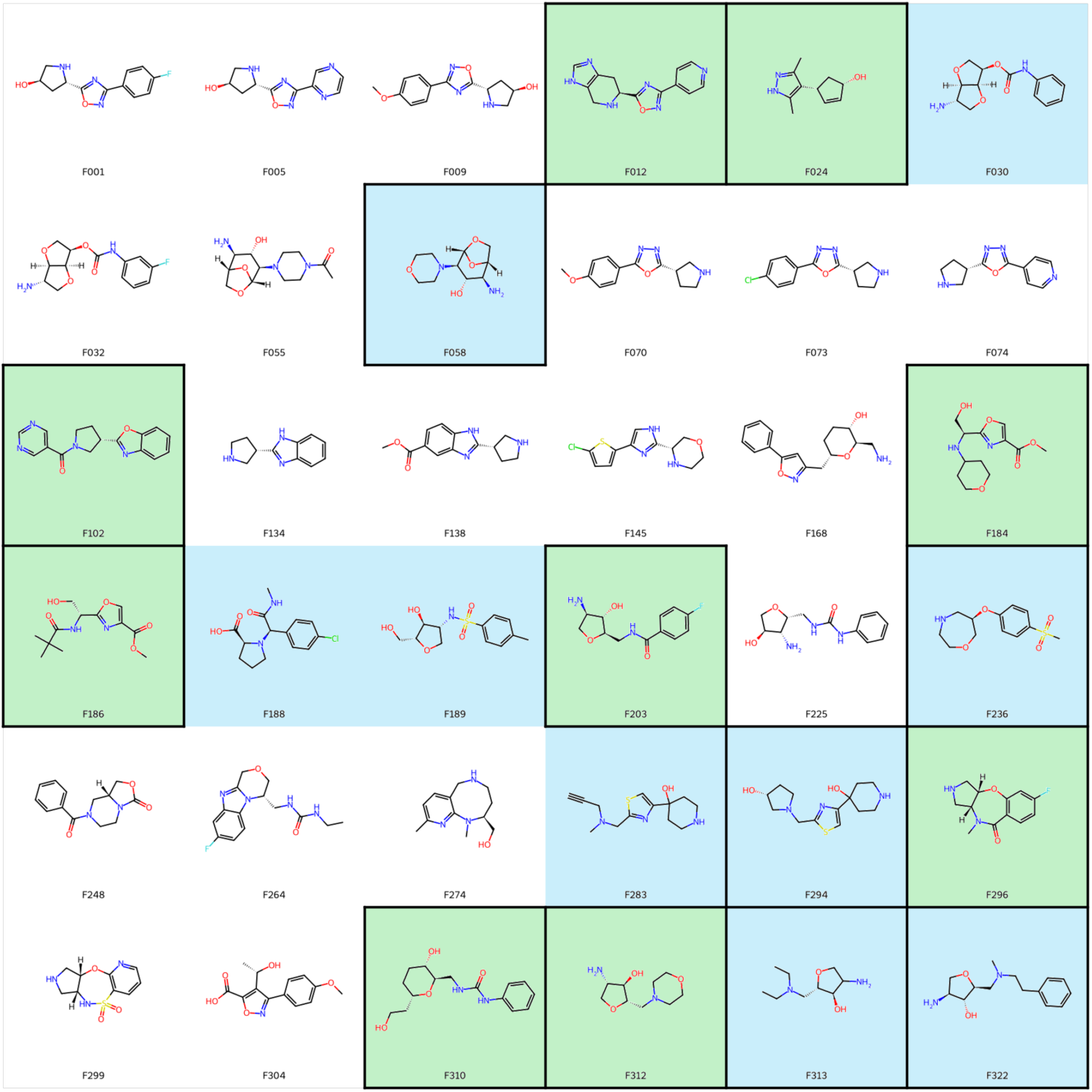
Chemical structures of the 36 fragment hits. Fragments that bind only in the ATP pocket are highlighted in green; fragments that bind only at one or more peripheral site(s) are highlighted in blue; fragments that bind to a single site are indicated by a black frame. All other fragments bind both in the ATP pocket and simultaneously to at least one additional peripheral site.

### 2.2 Prior Knowledge on the Fragments and their Natural Product Parents from Database Mining

First, we analyzed what was already known about the fragments detected as hits in the public domain, querying PubChem and ChEMBL.^42,43^ Although 35 fragments possess a PubChem entry, none of them is associated with any target or bioactivity annotation in this database. In ChEMBL four fragments are annotated, namely F012 (CHEMBL3437495), F074 (CHEMBL3437482), F186 (CHEMBL4937704) and F236 (CHEMBL4916673), whereby bioactivity data and target annotations are only available for the first two. For both, bioactivity data are reported for bacterial and protozoan protein targets only: methionyl-tRNA synthetase from *Leishmania donovani*; histidine-tRNA ligase from *Leishmania infantum* or *Trypanosoma cruzi*; ClpP1P2 protease, coenzyme A biosynthesis bifunctional protein, polyketide synthase and tryptophan-tRNA ligase from *Mycobacterium tuberculosis*; lysine-tRNA ligase from *Plasmodium falciparum*. Notably, no activity towards any kinase target has yet been reported.

As the fragments were deduced from NP, we investigated what is already known about NPs comprising the fragments as substructures, particularly with respect to their biological activities. A substructure search in the COlleCtion of Open NatUral producTs (COCONUT) database^44,45^ yielded 1675 different NPs. Depending on the individual fragment, the number of associated NPs largely varied from 0 to 554 (Fig. S2). No NP was found in the COCONUT for F001, F032, F203, F264, and F296, all of which are fluorinated. According to the COCONUT curators, many of such compounds have been removed from their database, as there is a lack of conclusive evidence supporting their classification as NPs.^45^ Unsurprisingly, all the non-fluorinated fragments were found to possess an entry in the COCONUT database themselves, as the latter contains molecules from the subset of NPs in the ZINC15 database, among others, which in turn integrates the AnalytiCon Discovery libraries. More than a hundred NPs are associated with F030, F313, and F134. In the following section, we will pinpoint potential growth vectors in these fragments that may be of possible interest for the follow-up development of the identified fragment hits.

Of the 1675 NPs found in COCONUT comprising the fragments as substructures, 1483 were found to possess a PubChem entry, and 93 were stored in ChEMBL. In PubChem, 82 target annotations were available for 54 NPs, covering 14 distinct protein targets. Of these, 70% are associated with receptor (-like) proteins, 15% with methyl transferases and 9% with lipoxygenases. Individual entries were identified for a hydrolase and an amino transferase, as well as for inositol hexakisphosphate kinase 1 (IP6K1). IP6K1 is a member of the inositol polyphosphate kinase (IPK) family, which catalyze the transfer of phosphoryl groups from ATP to various inositol phosphates. Although some structural homologies have been reported between kinases from IPK family (PFAM-ID: PF03770^46^) and the eukaryotic protein kinase (ePK) family, which includes PKA, they are only very distantly related with respect to their enzymatic properties.^47,48^ Hence, no assumptions on the NP’s bioactivity towards ePK or PKA can be drawn solely based on the reported IP6K1 bioactivity.

ChEMBL provides target annotations including bioactivity data with an assay confidence score of 9 – indicating direct testing of a single protein target – for 23 NPs, covering three distinct protein targets. Thereof, 87% of the NPs exhibit up to 50% inhibition of histidine-tRNA ligase from the protozoan organism *leishmania infantum*. Two NPs have been tested for their inhibitory activity against human glycogen synthase kinase-3 beta (GSK-3β) and one NP against P-glycoprotein (P-gp). However, all three NPs are inactive against GSK-3β and P-gp, respectively, with IC_50_ values above 50 µM.

Overall, little is known in the public domain about the bioactivity of the NPs that comprise the discovered fragments as substructures, or the fragments themselves. To date, none of the fragments or associated NPs have been reported to be active against any of the 518 human protein kinases with the conserved structure of the catalytic domain shared with PKA.

### 2.3 Growth Vector Analysis and Fragment Sociability

Many FBDD projects have been discontinued in the past due to synthetic intractability or the effort of chemistry required for structural expansion and optimization of the fragment hits.^21^ This is also why NPs in particular have received little attention as a pool for fragment-based screening earlier.^18^ However, this is expected not to be an issue in our case, as the concept of ‘fragment sociability’^21^ has already been integrated into the initial fragment design/selection. A fragment is considered as ‘social’ if a significant number of close analogs is commercially available and if robust synthetic methodology allows the elaboration of growth vectors.^21^

In this context, we analyzed the derivatization points/patterns of the natural product parents to reveal possible growth vectors in the fragments (Fig. 2 and S3). In addition, we will next address the aspect of the commercial availability of possible follow-up compounds.

**Figure 2:**
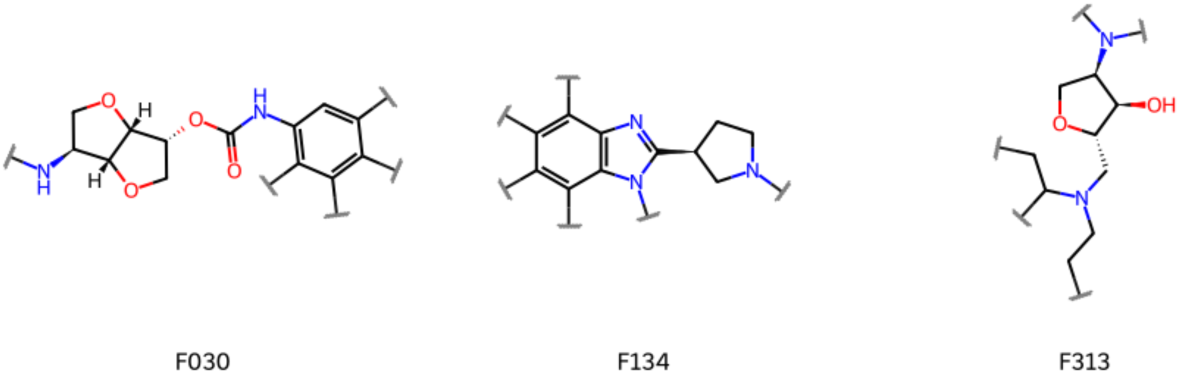
Growth vectors identified in the fragments with more than 100 associated natural product parents (F030, F134, and F313, see previous section), based on derivatization patterns of the latter. Growth vectors are indicated by squiggled lines perpendicular to a bond. Note that not all of them may be compatible with the binding mode evident from the complex crystal structure.

To quantify ‘fragment sociability’ we used the NATx library, which provides multiple follow-up compounds with exemplified growth vectors off-the-shelf, hence enabling efficient validation, examination of structure-activity relationships, and ultimately rapid optimization of the fragment hits (’growing by catalog’).^18,41,49^ On average, each fragment has 405 follow-up compounds sharing the same chemotype, 172 with the same scaffold, and 28 containing it as a substructure available from the NATx library. However, when selecting candidate molecules it must be considered that not all modifications may be compatible with the crystallographically determined binding mode in general, or may be favorably contributing to kinase binding/inhibition. Overall, the NATx library alone already provides a rich resource of NP-inspired molecules for follow-up studies.

### 2.4 Structure-Based Analyses

#### 2.4.1 Structural Classification of Protein Kinases and their Ligands

When describing a kinase and its role in ligand binding, two aspects are to be considered, the conformational state of the protein and the type of ligand in the complex.

All 36 PKA fragment complexes exhibit the DFG-in, αC-helix-in conformation, which is also the most frequently observed conformation for ligand-bound PKA structures in the PDB (99% of the 239 PKA:ligand complex structures used as reference herein). While the DFG-in (and DFG-out) label only provides a broad description of a more complex kinase conformational landscape, the more sophisticated Modi and Dunbrack scheme distinguishes 8 kinase conformations based on the dihedral angles of the xDFG motif residues. They are named after the region occupied in the Ramachandran plot (A = alpha; B = beta; L = left) and the phenylalanine rotamer (minus, plus, trans).^50,51^ According to this scheme, 6 subclasses of the DFG-in conformation were identified: BLAminus, BLBplus, ABAminus, BLBminus, BLBtrans, and BLAplus. 35 of the complex structures are in the so-called BLAminus conformation, corresponding to the kinase active form, while one occupies the active-like ABAminus conformation (F274). ABAminus structures strongly resemble the active BLAminus state, involving only a peptide flip of the x and D residues of the xDFG motif. In the remaining part, the positions of the activation loop and the αC-helix are very similar.^50,51^ Nevertheless, the ABAminus conformation corresponds to an inactive kinase state, as the aspartate does not facilitate the positioning of the cofactors ATP and Mg^2+^.^50^ These two conformational classes also correspond to the most common ones found for ligand-bound PKA structures in the PDB, with abundances of 80% and 18%, respectively.

Following the nomenclature introduced by Dar and Shokat^52^ and extended by Gavrin *et al.*,^53^ all 36 fragments correspond to type I and/or type IV ligands. Most of the fragments bind to multiple binding sites in the protein, most frequently in the ATP binding pocket and one of the peripheral sites. The fact that fragments tend to bind at several individual binding sites, has been frequently observed and is due to their smaller size and fewer options to establish interactions, as well as to the fact that fragments are usually added in higher concentrations to the crystallization conditions (*vide infra*).^27^ However, 9 fragments bind exclusively to peripheral sites and 9 are pure ATP site binders.

The fragment binding modes will be discussed in sections 2.4.3 – 2.4.4 and 2.5.1.

#### 2.4.2 Detailed Pocket Analysis

Next, we analyze in greater detail, which binding sites are occupied by our fragments and other known PKA ligands, respectively (Fig. S4). PDB ligands were found to occupy mostly the ATP pocket A, but also the peripheral pockets B, F, G and K, defined by Laufkötter *et al.*.^36^ Those bound fragments, which were successfully assigned to the Laufkötter pockets, bind to almost the same set of sites as the PDB ligands, namely A, E, F and G (Fig. 3). Site E is located in the C-terminal lobe, distal to the active site. To date, in no PDB structure of PKA a ligand is found to bind to this site, while it is occupied by our F189 (Fig. S6).

**Figure 3:**
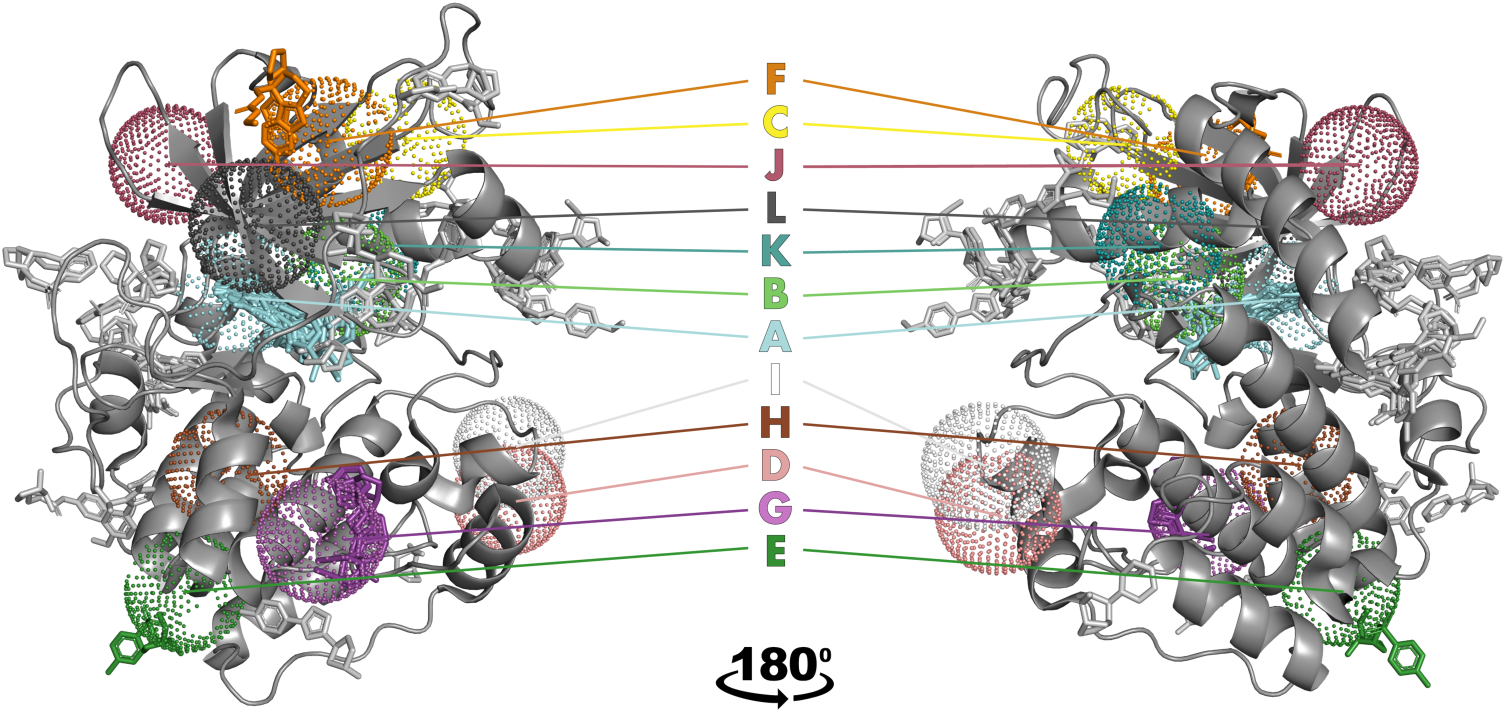
The kinase catalytic domain comprises 12 binding pockets in total (designated A–L), including the ATP site (A), and 11 additional binding pockets (B–L) as described by Laufkötter *et al*. ^36^ Pockets (in dot representation), defined by a 5 Å radius around the center of mass of all reference ligands (see Materials & Methods), are mapped on a PKA structure (PDB-ID: 3FJQ, cartoon representation). Note that according to Laufkötter *et al*. no individual kinase contains all the pockets.^36^ Fragments are colored the same way as the pockets they occupy (no element-specific coloration). Fragments that could not be assigned to any of the known kinase pockets (52.2% of all fragment copies) are displayed in light gray.

Site E is sometimes referred to as the ‘myristoyl pocket’ as myristic acid was the first ligand found to bind there in ABL1 kinase.^54^ While PKA membrane association and thus activity is also regulated via *N*-myristoylation, its myristoyl binding site is located in a different area (covalent linkage via Gly1, PDB-ID: 1CMK) than in ABL1 kinase.^55^ Site E has been reported to accommodate type IV inhibitors as well as activators, with the most prominent example being the approved PKI asciminib, targeting the BCR-ABL1 tyrosine kinase domain (Fig. S6).^38,56^ Asciminib reinforces the pharmacological relevance of allosteric kinase ligands and is an example of the general promise that allosteric inhibitors are likely to be more selective and resistant to mutation than ATP site binders as they target sites with low conservation throughout the kinome. Lastly, it should be mentioned that the C-terminal αI-helix of tyrosine kinases such as ABL1 exhibits significant conformational flexibility and borders the allosteric pocket E in its kinked conformation,^57^ which is not the case for PKA. Ligands binding to allosteric site E in kinases with a straight αI-helix, which does not enclose the pocket, have also been identified. However, only those that induce αI-helix bending function as allosteric inhibitors (in ABL1).^57^ Whether our fragment F189 can bind to the equivalent site in a pharmacologically relevant target such as ABL1 kinase and trigger the conformational change required for inhibition will be addressed in a subsequent study.

The other fragments assigned to a Laufkötter peripheral site are F134 and F264 (both binding to site F, the ATP site and at least one additional peripheral site), as well as F030, F138 and F283 (all binding to peripheral site G and at least one other site).

Interestingly, 85.7% of all non-ATP site binders could not be assigned to any of the kinase pockets reported by Laufkötter *et al.*,^36^ for which at least one ligand was demonstrated to modulate activity of at least one kinase via an allosteric mechanism. Since the bioactivity of these fragments was not the scope of our study, it is possible that some represent (surface-bound) fragments without functional relevance, rather than being true modulators of kinase activity. Moreover, the assignment may be hindered by imperfect superimposition of the PKA and the reference structures that originate from various kinases and different kinase families. Also, a general caveat of the crystallographic fragment screening approach should be borne in mind: Contact sites in the crystal packing are often detected as pockets or at spots and accommodate fragments. Such hits are typically specific to the crystal environment and will likely not bind to the protein in solution. In our study, this accounts for only a minor portion of the obtained hits which means 14 out of 69 (20.3%) of all fragment copies bind in proximity to more than one PKA molecule in the crystal lattice. Among them, 5 peripheral binders are unlikely to be crystal-packing related since *>*50% of their surface area is closer to the asymmetric unit than to the symmetry mate, and they form more polar interactions with the asymmetric unit than with the symmetry mate. Whether our fragments or derived lead compounds also bind and modulate a kinase in solution will be examined in a subsequent study.

Below, we will focus on describing the interaction of F189 with the two neighboring PKA molecules in the crystal lattice (Fig. S7) to verify its specific binding. The interactions of F189 within the asymmetric unit involve seven hydrogen bonds – to the residues His260, Glu140, and Arg144, as well as to three water molecules. In contrast, the interactions of F189 with the protein symmetry mate include only a hydrogen bond to a water molecule in the first hydration shell and a hydrophobic interaction of the phenyl ring with the hydrophobic pocket. Concerning the fragment’s surface area, however, both PKA macromolecules contribute almost equally to the binding site, with 54.1% and 45.4%. Despite the interactions of F189 with the symmetry mate, which of course, also favor the fragment binding at this site, it can be assumed that binding is indeed a specific binding event at a yet underexplored allosteric site.

Fragment F189 has 405 potential follow-up compounds sharing its chemotype commercially available from the NATx library alone, thereof 175 with the same scaffold, and 4 containing it as a substructure.

#### 2.4.3 Fragment Binding Modes

Figures 4 and S1 display the binding modes of all ATP site binders. Nearly half of them are hinge binders, forming one or two hydrogen bonds to the protein backbone of the hinge residues (Glu121, Tyr122, Val123), and/or featuring an ionic interaction with the side chain of Glu127, which is part of the linker region following the hinge. Furthermore, many of the ATP site binding fragments form a hydrogen bond and/or ionic interaction with the Thr183 and the Asp184 residues of the conserved xDFG motif; and/or form one or two hydrogen bonds/salt bridges with residues from the catalytic loop (Glu170, Asn171); and/or the glycine-rich loop (Leu49, Thr51, Phe54).

**Figure 4:**
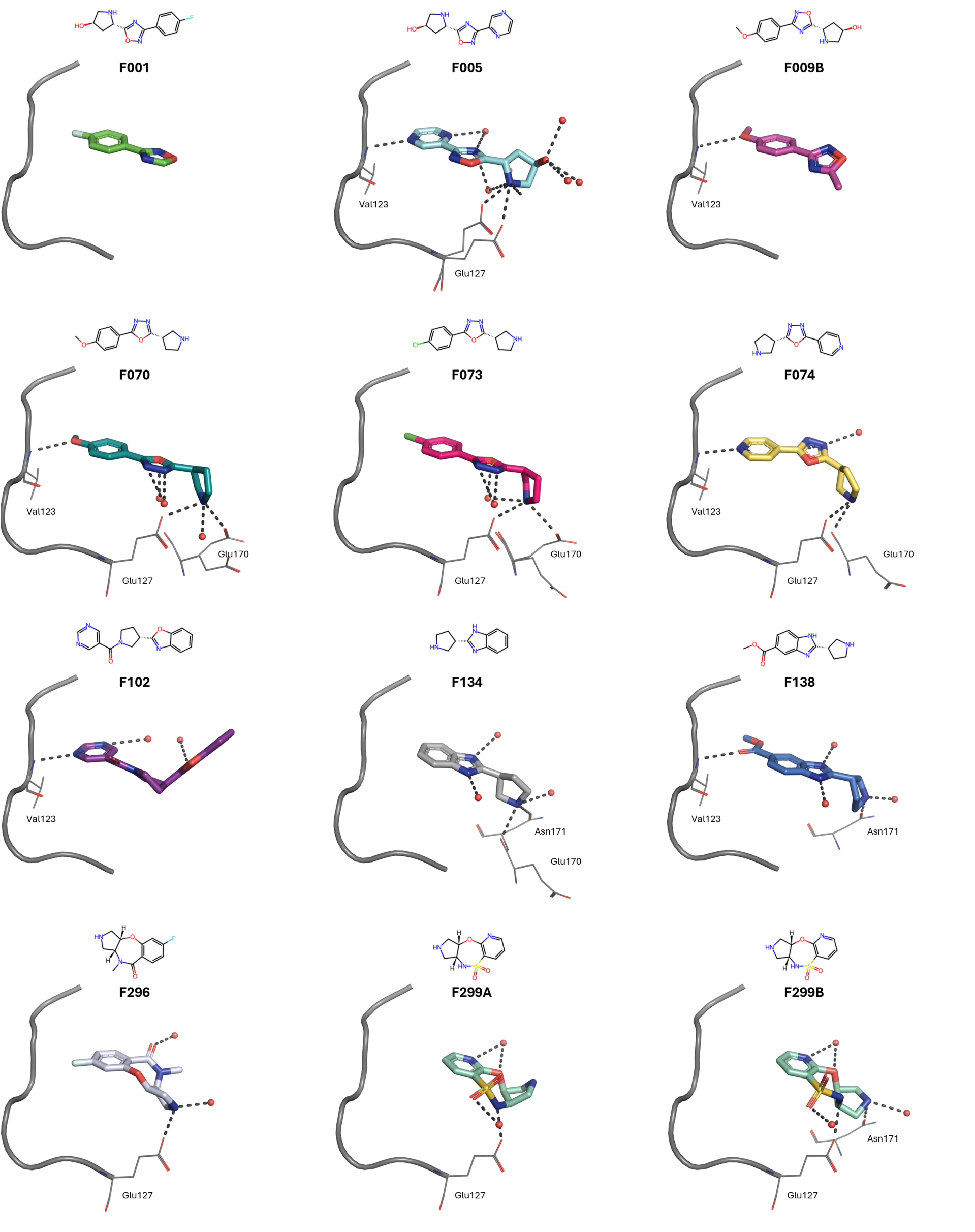
Binding modes of selected fragments within PKA’s ATP pocket. If multiple conformations of the fragments were resolved, we show each individually (A/B). Fragments are shown as sticks, amino acid residues involved in polar interactions as lines, waters involved in polar interactions as red spheres, and polar interactions as dashed lines. Additionally, the kinase hinge and linker (residues 120 – 127) are shown in cartoon representation. For clarity, only amino acid residues involved in polar interactions are labeled. More fragment binding mode depictions can be found in Fig. S1.

Some of the fragment hits possess the same scaffold/chemotype. Their binding modes will be compared in more detail in the following. Binding modes of fragments sharing their chemotypes with PKA ligands from the PDB will be described later on.

F001, F005 and F009 belong to the same chemotype, so do F070, F073, and F074. All of which bind within the ATP-binding site, with their 1,2,4-oxadiazole or 1,3,4-oxadiazole ring in roughly the same position, and a (substituted) 6-membered aromatic ring pointing towards the hinge region. The latter is involved in hydrogen/halogen bonding with the hinge residues in all cases, except for F001, which lacks a suitable functional group. On the contrary, the oxadiazole does not form a hydrogen bond to any amino acid residue, which is presumably why its orientation is not conserved. F134 and F138 share the same scaffold and bind in the same orientation within the ATP binding site. However, their exact binding location is slightly shifted, as the additional methyl ester group of F138 occupies the space in the proximity of the hinge residues. Its carbonyl oxygen is involved in a hydrogen bond to Val123. The positively charged pyrrolidine nitrogen of F134 forms a salt bridge to Glu170 as well as a hydrogen bond to Asn171, while only the latter is found for F138. Although F102 exhibits the same chemotype as F134 and F138, its orientation in the ATP binding site is inverted, so that the nitrogen of the additional pyrimidine ring builds a hydrogen bond with hinge residue Val123. F296 and the structurally similar F299 are also laterally displaced within the ATP binding pocket and form salt bridges with the aforementioned acidic residues. F203, F312, F313 and F322 all represent a 2-substituted (*2S,3R,4S*)-4-amino-3-hydroxy-tetrahydrofuran. While F203 and F312 bind within the ATP pocket in proximity to the hinge residues, and in the case of F312 forming a hydrogen bond with Val123, the other two fragments bind at the other end of the pocket, in the phosphate subpocket and beyond. Beyond the ATP site binders, F030 and F032 feature a conserved binding mode within peripheral pocket G (2.4.2 Detailed Pocket Analysis). On the contrary, the binding sites/modes of three structurally similar fragment sets (F055 + F058, F184 + F186, F283 + F294) are not conserved. While the binding modes are preserved for most of the structurally similar fragments, this is not the case for all of them.

#### 2.4.4 Unexpected Binding Modality of F274

As already discussed above, the complex structure with F274 occupies the active-like ABAminus conformation characterized by a flip of the peptide bond connecting the x (Thr183) and D (Asp184) residues of the xDFG motif relative to the active BLAminus kinase conformation. Consequently, the side chains of both residues also occupy alternative conformations, enabling the formation of a hydrogen bond between the Asp184 side chain and the ligand’s hydroxyl group (Fig. 5). With an occupancy of 0.4, the typical xDFG conformation of the kinase in the active state can also be observed in the electron density. As the structural adaptation is minor, it is plausible that this change could even be induced in a soaking experiment, where ligands are added to pre-formed apo protein crystals, which typically occupy the BLAminus conformation, as seen in PDB structures 4WIH and 6EH0.^50^

**Figure 5:**
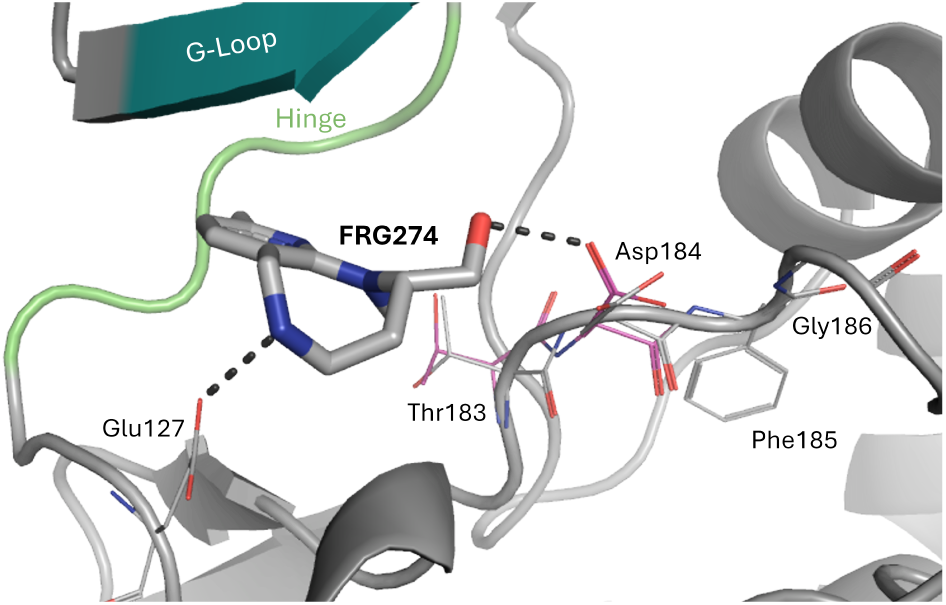
Unexpected binding modality of F274. The xDFG residues in the ABAminus conformation are displayed in color, while their alternative conformation is shown in gray. Hydrogen bonds are visualized as black dashed lines. For better clarity, water molecules are not displayed and the view differs from the common one.

It should be noted that Modi and Dunbrack^50^ have demonstrated with the help of the electron density score for individual atoms (EDIA)^58^ that in many cases ABAminus structures are in fact mismodelled BLAminus conformations. An analogous examination proves that this is not the case for our F274 complex structure nor for the structures discussed below (xDFG motif residues’ electron density score for multiple atoms (EDIA_m_) scores *→* 0.78, *i.e.*, well-covered by the electron density).

Even though the pharmacological relevance of the ABAminus conformation remains to be elucidated, the (presumably ligand-induced) peptide bond flip does not appear to prevent a ligand’s inhibitory effect and suitability to be developed into a PKI, as exemplified by multiple PKIs on the market. However, some kinase drug complex structures show inconsistent kinase conformations.^50^ For instance, the PDB contains complex structures of fragment-sized fasudil (M77) bound to the catalytic subunit α (PKACα) from *Cricetulus griseus* in both, the ABAminus (PDB-IDs: 5LCP, 5NW8, 5O0E, 5OK3, 6ERW, 6I2A, 6I2C) and the BLAminus conformation (PDB-IDs: 6EM2, 6YNA). In this example, a correlation between conformation and applied crystallization technique can be observed: ABAminus structures were obtained by co-crystallization, while BLAminus conformations originated from soaking experiments. Directly connected to this and even more critical with regard to the putative conformation change, is the ligand occupancy: In the soaked structures, the occupancy is below 1, whereas it is 1 in the co-crystallized complexes. For F274, the fragment copy binding to the ATP site is likewise refined with full occupancy. In addition to the seven fasudil-bound ABAminus structures, 37 further PKACα complex structures occupying the ABAminus conformation are deposited in the PDB, all in complex with type I ligands (29 unique ligands). Of these, six structures were obtained by soaking (PDB-IDs: 3AMB, 3ZO2, 3ZO3, 4AXA, 4C37, 5N3D). Their ligand occupancies amounted to *→* 0.87. Altogether, these observations support the conclusion that ligands can indeed induce a conformational change from BLAminus to ABAminus in soaking experiments, but likely only if the ligand occupancy is high and they are sufficiently potent to induce the conformational transformation. Further commonalities among the ligands capable of inducing the conformational change remain to be elucidated.

#### 2.4.5 Conformational Analysis: Glycine-rich Loop (G-Loop)

Although known to undergo significant movement, the kinase G-loop conformation and its implication for ligand binding are only scarcely explored in kinase research.^59,60^ The G-loop is located above the ATP binding site and contains the highly conserved GxGxxG motif (^50^GTGSFG^55^ in PKACα). The latter is also known as the nucleotide-binding motif in accordance with its native function. The flexibility of the G-loop is an essential but seemingly unpredictable determinant of kinase inhibitor binding strengths.^61^ For instance, as described by Möbitz *et al.*,^60^ a stacking down of the loop onto the ligand creates a more buried cavity that may contribute to tighter binding and thus higher ligand efficiency.

A qualitative analysis of the G-loop conformations occupied in our PKA fragment complexes shows that two conformations stand out in particular, the one occupied with F055 and that with F322 (Fig. 6), in which the G-loop is stacked down onto the ligand, although neither interacts directly with the G-loop residues. On the contrary, F236 forms hydrogen bonds to the backbone of the G-loop residues Thr51, Ser53, and Phe54, while F186 interacts with the preceding and following residues Leu49 and Val57 (Fig. S1). In both complex structures, the common G-loop conformation is occupied. Owing to its high flexibility, the G-loop may not be fully resolved in X-ray crystallographic structures, as it is the case for the complexes with F134, F138, and F274. In other words, binding of these three fragments does not appear to stabilize a distinct G-loop conformation, as it appears for the other complex structures, possibly due to lower affinity or shorter residence time. Overall, the crystallographic data shown alone cannot provide any information about a putative coupling between the G-loop conformation and ligand binding. We plan to investigate this further in follow-up studies, using advanced methodologies such as molecular dynamics simulations.

**Figure 6:**
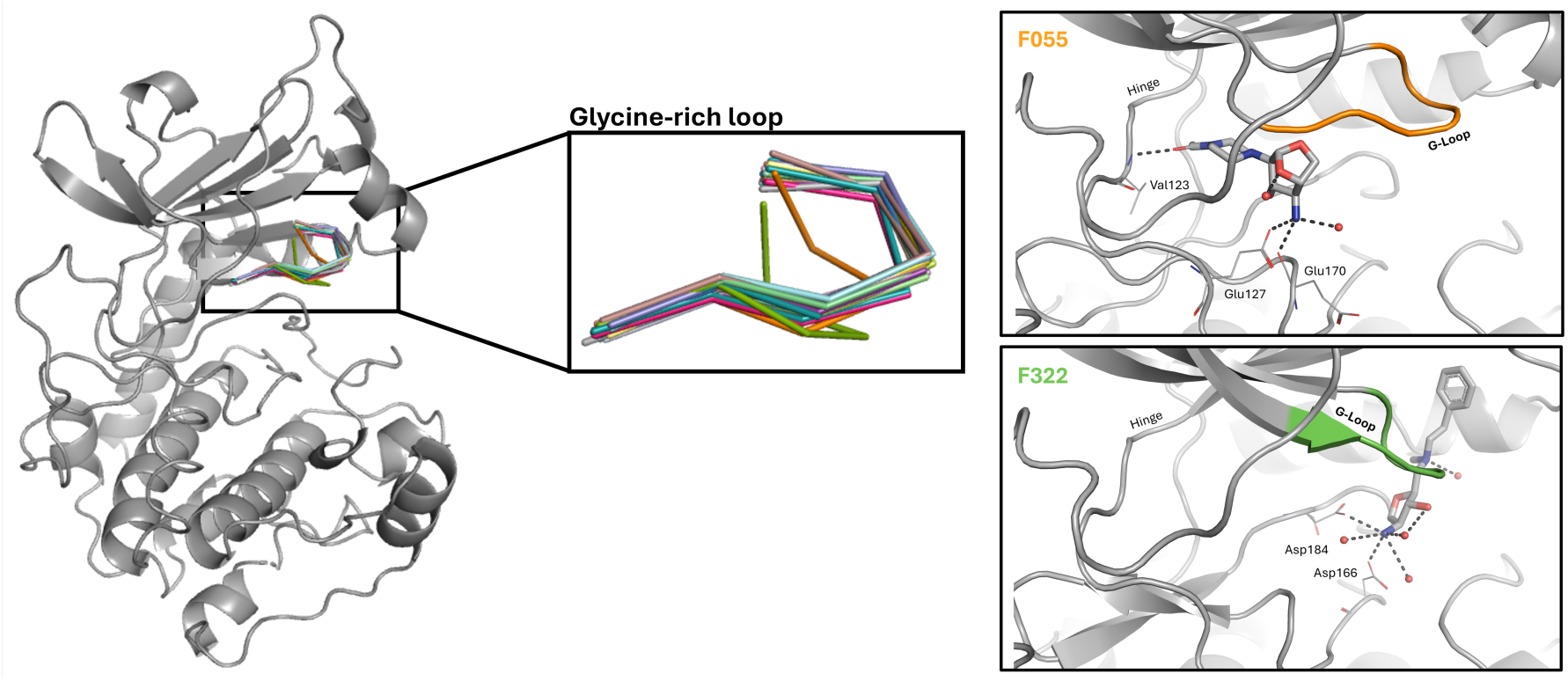
Superimposition of the glycine-rich loop (amino acid residues ^50^GTGSFG^55^, ribbon representation) of all our crystal structures. Note that in case of the complexes with F134, F138 and F274 not all G-loop residues have been modeled due to missing electron density support. The boxes on the right show the complex structures for which the G-loop conformation stood out, as it is stacked down onto the ligand. Therein, the G-loop residues are highlighted in the same color as in the left part of the figure. The protein is shown in cartoon representation and fragments are shown as sticks. Additionally, we show amino acid residues involved in polar interactions as lines, waters involved in polar interactions as red spheres, and polar interactions as black dashed lines.

### 2.5 Chemical Space Analysis

#### 2.5.1 Identification of Novel Scaffolds/Chemotypes

To assess the novelty of our fragment hits beyond the systematic database mining and the NPcentric analysis we compared their scaffolds across five reference datasets. In brief, these reference datasets correspond to two collections of compounds that bind PKA, extracted from the PDB and the ChEMBL database, respectively. Furthermore, we analyzed the data with respect to: A collection of compounds bioactive towards any typical kinase, as found in the ChEMBL database. A dataset encompassing the 107 protein kinase inhibitors (PKIs) currently on the market and a dataset of all approved oral drugs regardless of their biological target.

The 36 fragments represent 32 different Bemis-Murcko scaffolds, none of which have been previously reported in the reference datasets described above (Fig. S8). Among the 36 fragments, 21 different cyclic skeletons are included, of which two are not found in any of the reference datasets (Fig. 7), and 13 are absent from the PKA-specific reference datasets (Fig. S9).

**Figure 7:**
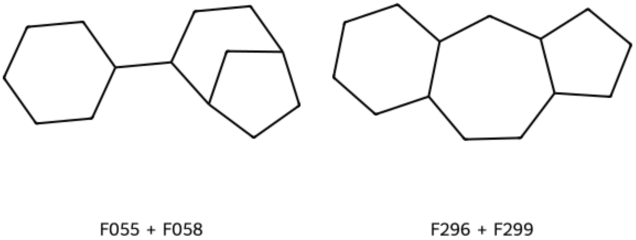
Cyclic skeletons of our fragments not found in any of the reference datasets.

The Bemis-Murcko scaffolds are by definition sensitive to minor structural modifications in the core of the molecule. Hence, we additionally inspected the three in terms of the Tanimoto coefficient top scored most similar reference molecules per fragment, irrespective from which reference dataset they originated (Figs. 8 and S10). The majority of the most similar reference molecules are contained in the ChEMBL Kinases dataset, which is by far the largest of our reference datasets. A few are PDB ligands, approved oral drugs or approved PKIs. The overall highest Tanimoto similarity of 0.50 was determined for F304 and mofezolac, an approved oral non-steroidal anti-inflammatory drug that selectively inhibits cyclooxygenase (COX1). F304 and putative follow-up compounds may also inhibit COX1 and thus may cause undesirable side effects. F012 and the PDB ligand T7W (PDB-ID: 7BB0) exhibit the second highest Tanimoto similarity of 0.46. F012 is a substructure of T7W, but shows (*S*)-configuration, while T7W is the (*R*)-enantiomer. Three more PDB ligands appear in the list of most similar reference molecules (Fig. 8), namely 8KB (PDB-ID), 8PQ (PDB-ID: 5N7U), and 7QI (PDB-ID: 7PID). 8KB represents a substructure of F236 and is thus even smaller than our fragment. 8PQ is an analog of F024 with a bromine substituent in the 4-position of the pyrazole ring, compared to the larger cyclopent-2-en-1-ol moiety in F024. 7QI and F145 share only a small proportion of their structure, namely the morpholine ring (linked to a biaryl). The binding modes of the fragment-PDB ligand pairs of the same chemotype are compared in the supporting information (Fig. S11). For approved PKIs, the highest Tanimoto similarity to a fragment amounts to 0.29, albeit largely differing in the size of the central ring, and in the type of N-heterocycle incorporated.

**Figure 8:**
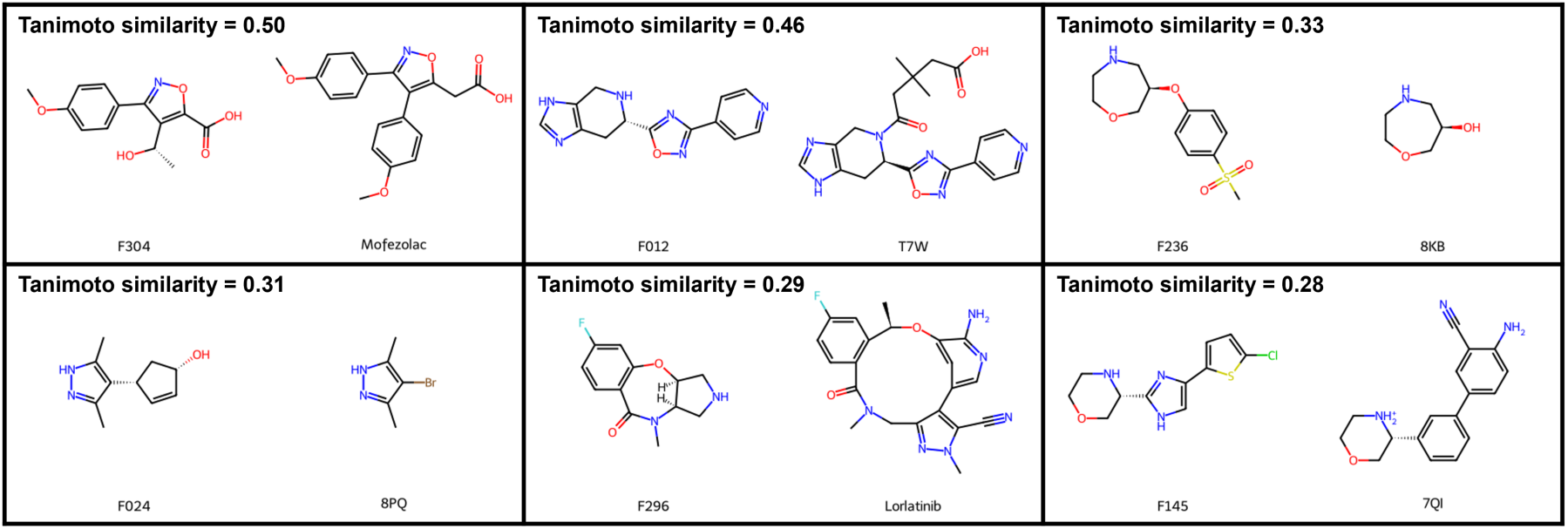
2D Structures of the in terms of the Tanimoto coefficient most similar reference ligands for the reference dataset of oral drugs, PDB ligands and approved PKIs, along with the related fragments. These also include the pairs with the overall highest Tanimoto similarity.

Although we have seen a few similar molecules in the reference datasets, this was only the case for a minority of the binders. Overall, the Tanimoto similarities are low, with a mean value of 0.31 for the most similar reference molecule per fragment. This again supports our argument that (most of) our fragment hits represent novel chemotypes for kinase targets. Altogether, the scaffold/chemotype analyses again highlight the novelty of our fragments, demonstrating their potential for further exploration, while generally avoiding conflicts in the already tightly occupied intellectual property space of kinase modulators, and enabling new insights into structural biology not observed before.

#### 2.5.2 Delineating the Fragment’s Chemical Space through Molecular Descriptors

To quantify the NP similarity of the fragments, their 3D character and physicochemical properties relevant to medicinal chemistry, and thus to delineate the occupied chemical space in comparison to the reference molecules, we analyzed various molecular descriptors (Fig. 9).

**Figure 9:**
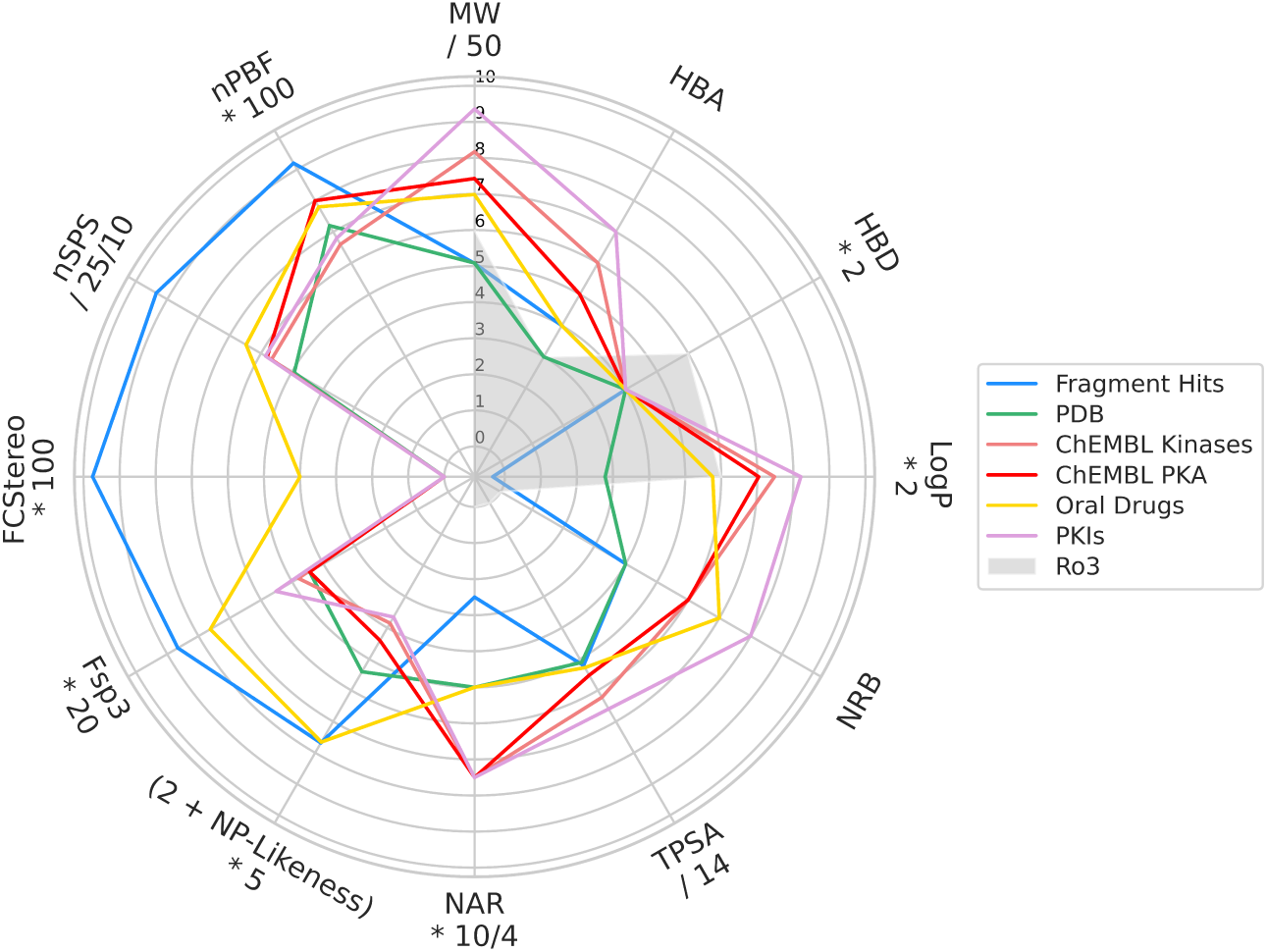
Median values of molecular descriptors. The descriptor space of the Rule of Three (Ro3)^62^ is indicated as gray shade. MW = Molecular weight. HBA and HBD = Number of hydrogen acceptor and donor atoms, respectively. LogP = calculated logarithm of the n-octanol/water partition coefficient. NRB = Number of rotatable bonds. TPSA = Topological polar surface area. NAR = Number of aromatic rings. NP likeness = Natural product likeness. Fsp^3^ = Fraction of sp^3^-hybridized carbons. FC_stereo_ = Fraction of stereogenic carbons. nSPS = Normalized spatial score. nPBF = Normalized deviation from the plane of best fit, calculated from *in silico*-generated ligand conformations. For detailed descriptions of the molecular descriptors refer to 5.4.3 Molecular Descriptors. Exact median values can be found in the supplementary data (Tab. S1). In addition to the median values, we present the descriptor value distributions and a tabular overview of the compounds/fragments associated with the highest/lowest descriptor value per dataset in the supporting information (Fig. S12 and Tab. S2).

Unsurprisingly, our fragments possess a lower molecular weight (MW), number of rotatable bonds (NRB), number of aromatic rings (NAR), lipophilicity (logP), and topological polar surface area (TPSA) compared to the molecules from the reference datasets described earlier. The average number of hydrogen bond donor (HBD) atoms is identical with a value of 2 across all data sets, including our fragments, despite their smaller molecular size. The average number of hydrogen bond acceptor (HBA) atoms spreads out over a larger range, whereby our fragments fall into the lower range compared to the reference datasets. Approved PKIs stand out, as they possess the highest median value of all seven aforementioned physicochemical descriptors, higher than that of the oral drugs. This is in concordance with the previous assessments in the literature^31,35^ and is presented here for bookkeeping purposes only.

For our fragment hits, the natural product (NP) likeness ranges from –1.82 to +0.88, with a median of –0.46. Lower median values are observed for the kinase-targeted reference datasets indicating that our fragments exhibit greater similarity to NPs than the reference molecules (all p-values *↑* 0.03, Fig. S12). In contrast, the NP likeness of the fragments is nearly identical to that of approved oral drugs. Interestingly, the NP likeness median value is lower for the NPs comprising the fragments as substructures (–0.97), compared to the one of the fragments themselves (Fig. S12). While the larger, more complex NPs include additional structural elements that may contribute to an increase in the NP likeness score, this is offset by the larger size of the NPs. In other words, the fragments exhibit a high(er) NP likeness score, as their structures are (more) densely packed with typical NP substructures than the parent NPs.

Looking at the fraction of sp^3^-hybridized carbons (Fsp^3^), the fraction of stereogenic carbons (FC_stereo_), and the normalized spatial score (nSPS) descriptors, our fragments possess a higher median value compared to the reference molecules. Dunn’s statistical test verified that the descriptor value distributions for the fragments differ significantly from those in the reference datasets with p-values *↑* 0.03. For the kinase-targeted reference datasets, all p-values are *↑* 10^−8^ (Fig. S12). In line with the findings by Lovering *et al.*^7^ who reported that the fraction of sp^3^-hybridized carbon atoms (Fsp^3^) measure correlates with overall clinical success, the datasets of approved PKIs and oral drugs feature higher Fsp^3^ median values compared to the datasets from ChEMBL and the PDB, which are dominated by early-stage discovery compounds (all p-values *↑* 0.004).

An outstanding proportion of 55.6% of our fragments surpasses the threshold value of Fsp^3^ *≥* 0.42 reported in the literature,^63^ while the percentages of compliant molecules in the kinase-targeted reference datasets amount to 16.2% at most (Fig. S13). The percentage of molecules surpassing the Fsp^3^ threshold of the dataset of oral drugs on the market is quite high (44.0%), again reflecting the finding by Lovering *et al.*.^7^ On the contrary, it amounts to only 15.9% for approved PKIs, mirroring the fact that most of them contain multiple aromatic rings which do not contribute to Fsp^3^, also indicated by the higher median number of aromatic rings in PKIs *versus* all approved oral drugs.

While all previously mentioned molecular descriptors were calculated from the molecule’s 2D structure, the deviation from the plane of best fit (PBF) descriptor was computed from the 3D structure, which was either generated *in silico* using RDKit’s ETKDG method^64,65^ or obtained from crystallographic data representing the protein-bound state. This two-legged approach was chosen to facilitate comparisons with reference datasets lacking 3D structural information, such as those derived from ChEMBL, while also leveraging available experimental structural data where possible. As the molecules differ largely in size across and within the datasets, the deviation from the plane of best fit descriptor has been normalized by the number of non-hydrogen atoms (normalized deviation from the plane of best fit (nPBF)).

One frequently criticized aspect in the field is the lack of attention to how molecular conformers are generated for descriptor calculation. To validate our approach, we compared (i) the *in silico*-generated and crystallographically determined conformations by means of the non-hydrogen atom root-mean-square deviation (RMSD) value, and (ii) the difference in the nPBF value calculated from both conformations. The median RMSD value amounted to 1.0 Å for the fragments and 1.5 Å for the PDB ligands. 95% of all fragment copies and 78% of all PDB ligand copies showed an RMSD *↑* 2.0 Å. This demonstrates the convincing relevance of the RDKit’s ETKDG-generated conformers,^64–66^ in particular for fragment-sized molecules. The largest difference in the nPBF value [ΔnPBF = nPBF(protein-bound) – nPBF(*in silico*)] was observed for PDB ligand GGB (5N3K), with ΔnPBF = –0.051 and RMSD = 0.4 Å. GGB is one of eight ligands for which no fully resolved copy was present in any crystal structure (see 5 Materials and Methods). The second-largest difference, however, was found for the fully resolved PDB ligand 46L (5N3E), with ΔnPBF = –0.039 and RMSD = 0.6 Å. Since only the largest ΔnPBF involves a truncated ligand and such cases make up a minuscule fraction of the dataset, their inclusion is not expected to bias the overall outcome, especially given that our analysis is mainly based on medians. We therefore retained these ligands to keep the datasets invariant and reflect real-world conditions in structure-based drug discovery, where partially resolved ligands are frequently encountered. Among the fragments, the largest difference was obtained for fully resolved F012 (ΔnPBF = –0.028, RMSD = 0.7 Å). Importantly, the nPBF values were not systematically higher or lower for the protein-bound *versus*the *in silico*-generated ligand conformations. Furthermore, no correlation was observed between RMSD and ΔnPBF values, as confirmed by Pearson and Spearman correlation analyses performed across both datasets.

With a median of 0.046, the nPBF value computed from the *in silico*-generated conformations is higher for the fragments compared to all reference datasets (Figs. 9 and S12). Dunn’s statistical test confirmed that the descriptor value distribution for the fragments differs significantly from those of the reference datasets, with p-values *≤* 0.02. Comparison with the kinase-targeted reference datasets only yields p-values *̤≤* 8×10^−4^ (Fig. S12). Calculation of the nPBF descriptor from the protein-bound ligand conformations was only applicable to the fragments and the PDB reference dataset. Like the nPBF values computed from the *in silico*-generated conformers, the distribution is shifted towards higher values for the fragments compared to PDB ligands (medians = 0.037 and 0.032; p-value = 5×10^−4^). Altogether, leveraging both crystallographically determined and *in silico*-generated ligand conformations enhances the reliability of our nPBF calculations, offering a more comprehensive view of molecular shape across datasets. We observe consistent trends for both approaches, with the fragments exhibiting higher nPBF values than the reference dataset(s) on average.

So far, the descriptor analyses have been binding site agnostic. However, for the fragments as well as for the PDB reference dataset, the binding sites have been classified earlier (2.4.2 Detailed Pocket Analysis). While allosteric inhibitors such as asciminib are gaining attention in kinase research, most PKIs continue to target the conserved ATP site. Ligands binding to this site typically comprise rather planar (sub)structures, including at least two aromatic rings.^30,35^ To explore whether the trends observed across the full dataset also apply to this key subset, we carried out a focused analysis of the ATP site ligands, encompassing 27 unique fragments and 151 distinct PDB ligands.

Indeed, we found that the ATP site-occupying fragments show the same statistically significant, favorable trends in their property distributions (Figs. 10, S12 and S13, Tab. S1). Also, the percentage of molecules exceeding the threshold of Fsp^3^ *≥* 0.42 is much higher for the ATP site fragments (40.7%) compared to the PDB ligands occupying the same site (13.9%).

**Figure 10:**
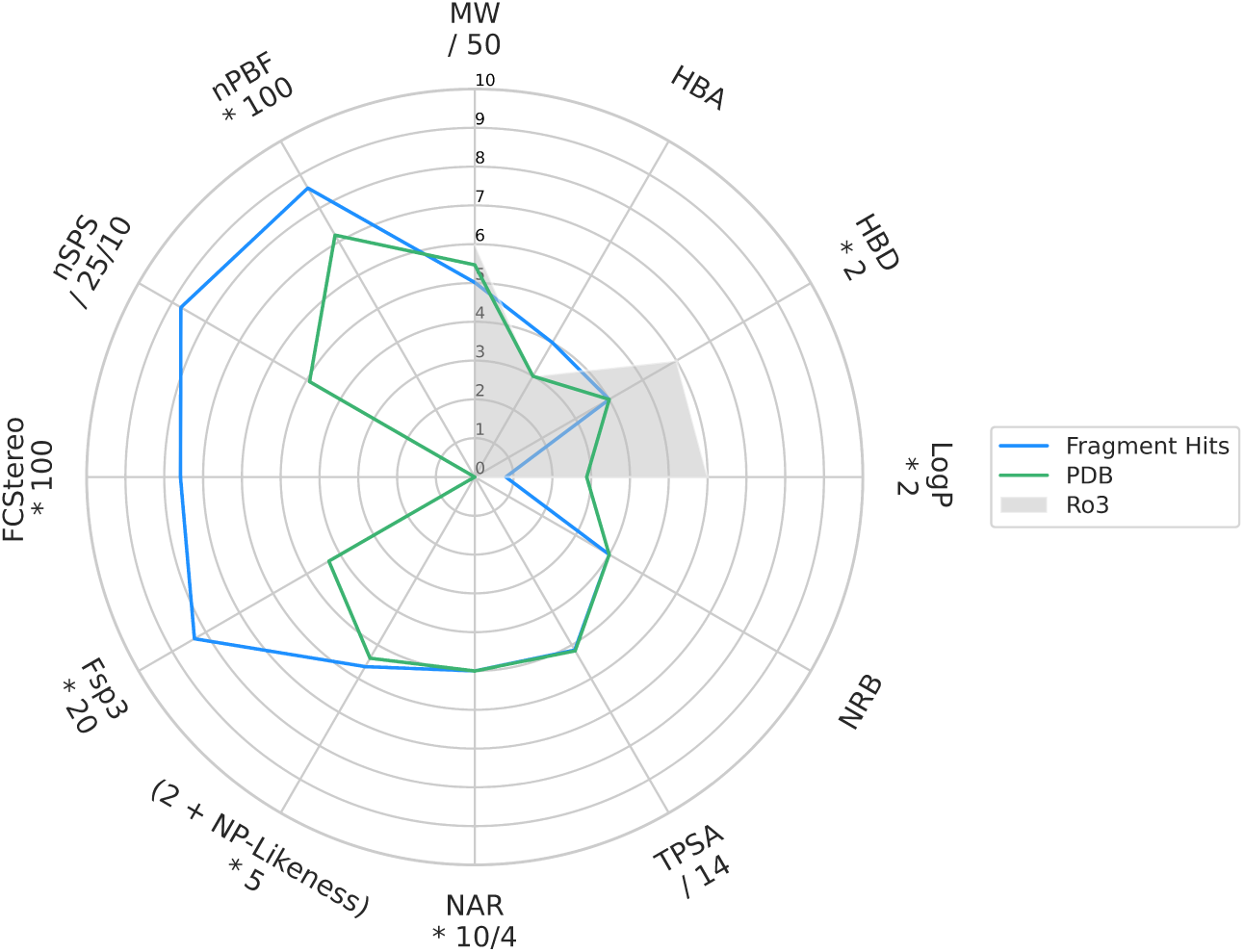
Median values of molecular descriptors for the subset of ATP site binders. The descriptor space of the Rule of Three (Ro3)^62^ is indicated as gray shade. MW = Molecular weight. HBA and HBD = Number of hydrogen acceptor and donor atoms, respectively. LogP = calculated logarithm of the n-octanol/water partition coefficient. NRB = Number of rotatable bonds. TPSA = Topological polar surface area. NAR = Number of aromatic rings. NP likeness = Natural product likeness. Fsp^3^ = Fraction of sp^3^-hybridized carbons. FC_stereo_ = Fraction of stereogenic carbons. nSPS = Normalized spatial score. nPBF = Normalized deviation from the plane of best fit, calculated from *in silico*-generated ligand conformations. For detailed descriptions of the molecular descriptors refer to 5.4.3 Molecular Descriptors. Exact median values can be found in Tab. S1. In addition to the median values, we present the descriptor value distributions in Fig. S12.

Taken together, the descriptor analysis revealed that our fragment hits exhibit above-average molecular three-dimensionality and spatial complexity, reduced aromaticity, as well as a higher degree in saturation compared to known kinase binders. This also holds true when looking at the subset of ATP site binders alone. Although the descriptor distributions for our fragments are statistically distinct from those of the reference sets, there is considerable overlap. In other words, even within the reference sets, some molecules also exhibit high values of these descriptors. Thus, our fragment hits rather expand than redefine the accessible chemical space for kinase binders by populating less frequently sampled regions. Nevertheless, the different distribution patterns highlight that our library of natural product-like fragments is tailored towards improved three-dimensional properties, while preserving a high hit rate.

Finally, we take a closer look at the individual descriptor values obtained for our fragment hits. In the case of Fsp^3^, FC_stereo_, and nSPS the five highest descriptor values are associated with the same set of fragments (F055, F058, F312, F313), albeit in diverging order (Tab. S2). An exception to this is the nPBF descriptor, for which F312 and F313 score the highest. Additionally, F024 and F189 or F058 and F264 are among the top-scoring fragments depending on whether the nPBF descriptor has been computed from the *in silico*-generated conformation or the protein-bound one. Interestingly, F313, F058, F312, F322, and F055 also exhibit the highest NP likeness values of all fragments (in decreasing order). This reflects the general trend that natural products, unlike many synthetic compounds, typically possess a higher degree of saturation and more stereocenters^14^ – a characteristic also captured by our NP-like fragment hits.

**Table 1:**
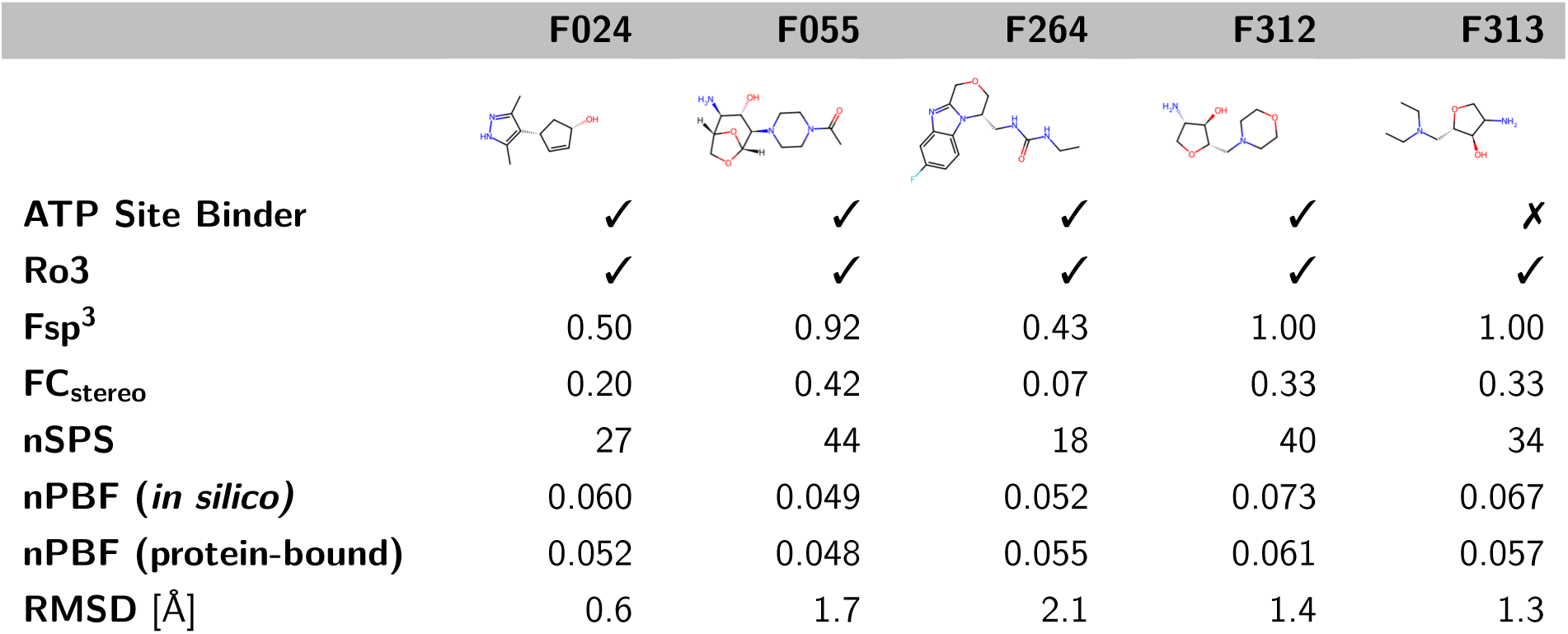
Exemplary selection of fragments, along with their descriptor values and Rule of Three compliance. The RMSD value denotes the mean structural deviation between the fragment’s *in silico*–generated conformer and all its observed conformations in the protein-bound state. Similarly, the nPBF value (protein-bound) represents a mean value across all crystallographically determined conformations.

|  | F024 | F055 | F264 | F312 | F313 |
| --- | --- | --- | --- | --- | --- |
| **ATP Site Binder** | ✓ | ✓ | ✓ | ✓ | ✗ |
| **Ro3** | ✓ | ✓ | ✓ | ✓ | ✓ |
| **Fsp<sup>3</sup>** | 0.50 | 0.92 | 0.43 | 1.00 | 1.00 |
| **FC<sub>stereo</sub>** | 0.20 | 0.42 | 0.07 | 0.33 | 0.33 |
| **nSPS** | 27 | 44 | 18 | 40 | 34 |
| **nPBF (*in silico*)** | 0.060 | 0.049 | 0.052 | 0.073 | 0.067 |
| **nPBF (protein-bound)** | 0.052 | 0.048 | 0.055 | 0.061 | 0.057 |
| **RMSD [Å]** | 0.6 | 1.7 | 2.1 | 1.4 | 1.3 |

Recently, Bournez *et al.*^35^ forecasted changes in the PKI landscape/chemical space in general, and new trends in terms of structures and molecular shapes in particular, based on their analysis of PKIs in clinical trials. Indeed, our analysis confirms this trend, as all the approved PKI for which the highest molecular descriptors for saturation, spatial complexity and molecular three-dimensionality were reported, namely gilteritinib (2018), peficitinib (2019) and repotrectinib (2023), have all been approved by the regulatory authorities in the last few years. We believe that the potential of such molecules as kinase modulators is not yet fully utilized and that these factors should receive greater attention in the rational design of such molecules, even in light of the recently published re-evaluation of the 15-year-old ‘*Escape from Flatland* ‘ paper.^7^ Churcher *et al.*^13^ did not find a clear relationship between the highest phase reached and Fsp^3^ after 2009 anymore. This apparent disconnect in the last years was attributed to the focus on kinases and the widespread usage of metal-catalyzed cross-coupling reactions, which made it easier to introduce sp2-hybridized systems into drugs.^13^ Nonetheless, our results suggest that exploring more saturated and three-dimensional chemical space holds promise for diversifying and advancing next-generation PKIs.

## 3 Conclusions

This study presented a crystal soaking campaign using 87 NP-like fragments against PKA, achieving an exceptionally high hit rate of 41%. These fragments are expected to be enriched in sp^3^-hybridized/ chiral carbons compared to conventional fragment libraries which usually exhibit a preponderance of sp2-hybridized carbons, aromatic and heterocyclic rings. This disproves the initial concerns that enhanced molecular 3D character would have a negative impact on the hit identification success.^2,10,11^ None of the fragments detected as hits, nor the natural products comprising them as substructures, have previously been reported as active against a typical protein kinase with the conserved catalytic domain shared by PKA in publicly available data. The 36 determined fragment-bound crystal structures, have been fully refined and represent a high-quality data set for compuational analyses. 35 structures displayed the kinase in the active (BLAminus) conformation, while the kinase in complex with F274 occupied the active-like (ABAminus) conformation. The latter was characterized by an unexpected fragment-induced flip of the peptide bond connecting the x (Thr183) and D (Asp184) residues of the xDFG motif. Out of the fragment hits, 9 corresponded to purely peripheral binders, 9 exclusively bound within the ATP site, and 18 fragments occupy simultaneously both orthosteric and peripheral sites. Notably, F189 was found to bind specifically at the underexplored allosteric site E, which is yet not occupied by any known PKA ligand stored in the PDB, but whose pharmacological relevance has been exemplified by asciminib (in BCR-ABL1 tyrosine kinase). A comparison of the fragments’ Bemis-Murcko scaffolds to the ones from five reference datasets, as well as examination of the most similar reference molecules further highlights the novelty of the fragment binders demonstrating their potential for further exploration, while avoiding conflicts in the already crowded intellectual property space of kinase binders, and enabling insights into structural biology not observed before. Further cheminformatics analysis revealed that these novel binders indeed populate less frequently sampled regions of chemical space characterized by enriched saturation, (spatial) complexity, and molecular 3D character. Specifically, we demonstrated that the fragments bound to PKA exhibit statistically significant shifts towards higher values in the Fsp^3^, FC_stereo_, nSPS, and nPBF descriptors compared to established kinase binders from reference datasets. The latter was computed from *in silico*-generated as well as from the crystallographically determined protein-bound ligand conformations, thereby enhancing the robustness of the shape analysis and ensuring comparability across datasets with and without experimentally resolved 3D structures. Notably, these favorable trends persist even within the subset of ATP site binders. These findings collectively emphasize how the resolved fragment hits challenge the planarity typical for kinase binders and venture beyond the flatland. While the spatial complexity is already inherent to the fragment’s scaffold, it can possibly be further enhanced through peripheral decorations in the hit or lead optimization phase. Rather than relying on tedious incremental growth and saturation of sp2-carbon-rich hits into 3D-shaped molecules by sp^3^-carbon chemistry, we already start with 3D-rich kinase-binding fragments, which can be expanded by established chemistry. In conclusion, the fragment hits identified using a high-performance soaking system for PKA present novel perspective on hit identification and lead optimization in kinase drug discovery.

## 4 Outlook

Huschmann *et al.*^18^ recently highlighted the utility of the FRGx library,^41^ a commercially available library of natural product-like fragments, for initial crystallographic screening and subsequent hit validation using readily accessible follow-up compounds. Our study supports this concept by showing that fragment hits from such a library populate less frequently sampled regions of kinase binder chemical space, characterized by an increase in saturation, sp^3^-hybridized carbons, stereo centers, and molecular 3D character. As the fragments themselves, we expect the potential follow-up molecules to possess the same favorable properties with all the mentioned benefits, including a higher likelihood of progressing to the next phase in the drug development process.

Beyond the development of traditional small-molecule inhibitors, novel kinase binders offer promising avenues for the design of proteolysis-targeting chimeras (PROTACs) – bifunctional molecules that induce selective proteasomal degradation of disease-relevant target proteins by recruiting them to E3 ubiquitin ligases. Given the high hit rate and flexible derivatization potential of NP-derived fragments, fragment-based approaches like ours could accelerate the development of PROTACs targeting kinases for which classical inhibition has proven insufficient or prone to resistance. Their defined 3D shape may confer scaffold rigidity and provide alternative exit vectors, allowing optimal spatial orientation of both ligase and target binding moieties and thereby facilitating the formation of the productive ternary complex (ligase–PROTAC–target). A prime example where PROTACs have shown promise in kinase research is Bruton’s tyrosine kinase (BTK), a critical target in B-cell malignancies such as mantle cell lymphoma and chronic lymphocytic leukemia.^67^ Several BTK-directed PROTACs, such as NX-2127 and BGB-16673, are already in clinical trials.^67^

In summary, the integration of natural product-inspired, three-dimensional fragments holds significant potential not only for the development of novel kinase inhibitors but also for advancing the design of modern drug modalities like PROTACs. This approach may be particularly beneficial in targeting kinases that are resistant to traditional inhibition strategies.

## 5 Materials and Methods

### 5.1 Fragment Selection

A selection of 87 fragments from AnalytiCon Discovery’s FRGx (’Fragments from Nature’) library^41^ was subjected to the soaking campaign. The selection was based on the availability and diversity of the fragments for the soaking experiments. In order to be consistent with the molecule identifiers used in the FRGx library, we herein stick with a discontinuous numbering of the fragment hits.

### 5.2 Crystallography

The full-length catalytic subunit alpha of PKA from Chinese hamster (*Cricetulus griseus*; UniProt-ID P25321, residues 1-350) was expressed, purified and crystallized as described by C. Siefker.^68^ Compared to the human enzyme, 6 amino acid replacements are given at positions 26, 35, 40, 43, 45 and 349, all of which distant (*>* 10 Å) from the ATP binding site, corresponding to a sequence identity of 98.3%. Fragments were dissolved in 100% DMSO to concentrations of 1 M or 500 mM. When fragments were not soluble at 500 mM, they were used as a saturated slurry. Soaking was performed by transferring one crystal each into a 2 µL drop of SmartSoak^69^-derived soaking condition which contained 10% of the fragment ligand stock. After 24 h of soaking, the crystals were harvested and flash-frozen. Datasets were collected at BESSY BL14.2 in Berlin.^70^ The required number of images, degree of rotation per image, and exposition time were calculated by taking test images before and after a 90° rotation and following the proposed strategy as computed by Mosflm.^71^ Data indexing, scaling, and integration were performed with XDS.^72^ Afterwards, a molecular replacement using an in-house structure of the PKA catalytic subunit was performed using PHASER.^73^ The structures were refined by iterative cycles with Phenix,^74^ while model building was performed with Coot.^75^ Sidechains which were not covered by electron density in the native maps were truncated to minimize model bias. Crystallographic tables can be found in the Supporting Information.

### 5.3 Analysis of Ligand-Bound Protein Kinase A Crystal Structures

Crystal structures were analyzed using Schrödinger’s PyMOL (version 2.6.0)^76^ together with home-written Python scripts. All structures were aligned to an ATP-bound PKA structure, 3FJQ. To put the results of the analysis of the novel 36 crystal structures into context, we further analyzed all PKA complex structures in the PDB (*vide infra*).

Pocket occupation of a ligand/fragment was defined as ‘being located in a 5 Å radius of an added and well-considered pseudo atom.’ In the case of the orthosteric ATP-pocket, the pseudo atom was placed at the center of mass of the adenine substructure of ATP, to capture all hinge binders. In the case of the allosteric kinase pockets, the reference structures as listed by Xerxa *et al.*^37^ were exploited, and the pseudo atom was placed in the center of mass of all ligands described to occupy the respective pocket.

To analyze if the binding of (peripheral) ligands may be influenced by the crystal environment we generated the symmetry mates within a cut-off distance of 6 Å of each ligand copy. Next, we determined if any ligand atom is closer to a neighboring symmetry mate in the crystal lattice than to the asymmetric unit itself. If so, we visualized the interactions of the ligand with each mate and computed the percentage of the ligand’s surface area closer to the respective mate.

For the structural classification of PKA and the fragment hits we utilized the KinCore^51^ standalone program (available from github) embedded in a bash script for batch processing of .pdb files.

Additionally, we evaluated the electron density support for the individual atoms in the FRG274:PKA complex structure, using the ediascorer 1.1.0 command line version. As we were primarily interested in the ABAminus conformation (occupancy = 0.6), we only kept this conformation of the residues Thr183 and Asp184 and manually removed the other from the .pdb file using a text editor. The electron density score for individual atoms (EDIA)^58^ quantifies the electron density support of each atom in a crystallographically resolved structure. An EDIA value *≥* 0.8 indicates that the atom is well-covered with electron density. Values between 0.4 and 0.8 or values *<* 0.4 point out minor or substantial inconsistencies with the electron density fit, respectively. Multiple EDIA values can be combined with the help of the power mean to compute EDIA_m_, the electron density score for multiple atoms to score a set of atoms such as a residue or a ligand.

### 5.4 Cheminformatic Analyses

The cheminformatics data analysis was performed using Python 3.10.13, together with Jupyter-Lab 8.6.0. Python script preparation and data analysis were done in Visual Studio Code 1.99.3 using a virtual environment managed in Mamba 1.5.8, with code version control via Git 2.34.1. Data retrieval through APIs was conducted using the requests 2.31.0 library for general HTTP requests, e. g. for querying COCONUT, KLIFS and PubChem, the biotite.database.rcsb module from biotite 0.39.0 for accessing PDB data, and the chembl_webresource_client 0.10.9 for retrieving information from the ChEMBL database. Data processing and analysis was performed with the help of Pandas 2.1.4, and NumPy 1.26.4 packages. Chemical data processing, including molecular descriptor calculation, substructure search, and scaffold generation, was conducted with RDKit 2024.03.5.^77^ Presented plots were prepared with matplotlib 3.8.3, seaborn 0.11.0 or ptitprince 0.2.7. All JupyterNotebooks and the .yml environment file are available on GitHub (https://github.com/czodrowskilab/NPFrag2Kinase).

#### 5.4.1 Reference Datasets

##### Natural Products

The COlleCtion of Open NatUral producTs (COCONUT) is one of the biggest and best annotated open-source databases for NPs.^44,45^ Canonical SMILES of the 695 133 NPs were downloaded from the COCONUT, version August 2024 (https://coconut.naturalproducts.net/download/). A stereospecific substructure search was conducted to identify all those NPs, comprising one of the fragments confirmed crystallographically as bound hits in this study as a substructure. Using the NPs’ SMILES as input, we queried two additional databases, PubChem^42^ and ChEMBL,^43,78^ via their APIs, for additional information on the molecules, including biological target annotations and bioactivity values.

##### Kinase-Targeted Reference Datasets

For the kinase-specific reference datasets, data mining was primarily based on the name or ID of the protein family (Pfam)^46^ of typical kinases that share the same fold as PKA, i. e., Protein kinase domain or PF00069. For PKA-specific datasets, filtering was refined using PKA’s enzyme commission (EC) number, being 2.7.11.11, along with limitation to the *α*-isoform of PKA’s catalytic subunit (PKACα).

ChEMBL (https://www.ebi.ac.uk/chembl/, version 34)^78^ is a curated database of more than two million molecules and their activities towards biological targets. First, we retrieved the target IDs of single proteins belonging to the protein kinase family, with protein classification ‘enzyme’ and ‘kinase’. In a subsequent step, the associated small molecules and bioactivity data were retrieved. After filtering for compounds that exhibit a pChEMBL value of *≥* 5.0 and/or a ligand efficiency^79^ of *≥* 0.3 towards any kinase in an assay with a confidence score of 9, and molecule standardization (*vide infra*), a total of 38721 compounds was obtained.

From the dataset described before, we created a subset of molecules bioactive towards PKACα from different species. Applied filters were PKA’s enzyme commission number (EC = 2.7.11.11), as it is non-species-specific, as well as the subunit and isoform indicators ‘catalytic’ and ‘alpha’ in the target name. In total, 71 molecules were identified to possess a pChEMBL value *≥* 5.0 and/or a LE *→* 0.3, towards PKACα, originating from three different species, namely Human (*Homo sapiens*), European Rabbit (*Oryctolagus cuniculus*) and Brown Rat (*Rattus norvegicus*), all of which possessing *>* 96% sequence identity.

The Protein Data Bank (PDB, https://www.rcsb.org/)^29^ collects 3D-structural data of biological macromolecules. First, the IDs of all structures tagged with the kinase domain Pfam name and the EC number of PKA were retrieved. Next, we used a GraphQL-based approach to retrieve the metadata associated with the structures. Structures corresponding to or comprising the regulatory subunit as well as complexes with protein substrates (of up to similar size as the PKA catalytic subunit), were excluded based on gene names. Subsequently, information on the non-polymer entities, including their molecular representation as SMILES, were extracted and uncomplexed structures were eliminated. As not all extracted molecules are ligands from a pharmacological point of view, but for instance simply originate as additives from the crystallization process, these should also be excluded from the analysis. Hence, we first filtered out all molecules that were included in the USAN Council’s list of pharmacological salts, regardless of charge status (e. g. acetic acid or acetate) and stereochemistry.^80^ In addition, other glycols/diols, sugars, fatty acids/lipids/lipid analogs/detergents, amino acids, solvents, and biochemical buffer molecules have been removed. As a result, no more than one ligand per PDB structure is considered in the analysis, yielding a total of 160 unique ligands, that are next subjected to molecular standardization (*vide infra*). It must be noted that some of these ligands have been co-crystallized multiple times, corresponding to a total of 239 PKA:ligand complex structures in the PDB, all of them determined by X-ray crystallography, with a median resolution of 1.8 Å. The protein component originates from five different species, namely Human (*Homo sapiens*), Chinese hamster (*Cricetulus griseus*), Cattle (*Bos taurus*), House mouse (*Mus musculus*), and Wild boar (*Sus scrofa*), all of which possessing *>* 97% sequence identity. The most recently deposited complex structure included in our analysis is 8SF8 (February 2024).

A dataset of approved drugs and clinical drugs targeting kinases is publicly available at https://klifs.net/drugs.php. This compilation integrates data from the Database of Protein Kinase Inhibitors in Clinical Trials (PKIDB)^31,35^ and the Kinase–Ligand Fingerprints and Structures (KLIFS) database.^32,81^ Therein included are 107 approved (clinical phase 4) PKIs. The most recently added PKI included in our analysis is Deuruxolitinib (July 2024).

##### Oral Drugs

In addition, we exploited ChEMBL as described earlier to compile a dataset of organic small molecules in clinical phase 4, *i.e.*, approved drugs that are administered orally. With the exclusion of prodrugs and compounds withdrawn from the market, and after molecule standardization (*vide infra*) and deduplication, the total number of drugs in this dataset amounts to 1064.

##### Data Preparation

While each of the mined databases have already implemented molecular standardization procedures, molecules for which no structural information was available were prepared using RDKit’s MolStandardize module to ensure consistency. This includes the disconnection of covalent bonds between metals and organic atoms (the disconnected metal is not preserved), desalting (keeping the largest fragment), neutralizing charges, SMILES canonicalization, and finally, deduplication. Alternatively, for those datasets, for which structural information was available, we used the protonation state(s) and tautomeric form(s) of the protein-bound ligand as determined by PROTOSS (*vide infra*).

Although few molecules were present in more than one of the datasets, none of the fragment hits has been found in the public datasets.

#### 5.4.2 Scaffold/Chemotype Analysis

Bemis and Murcko introduced the first formal classification of molecular scaffolds in 1996.^82^ According to their definition, a compound is composed of ring systems, linkers connecting the rings, and substituents. The compound’s scaffold is formed by the ring and linker atoms. Cyclic skeletons further generalize the Bemis-Murcko scaffolds, by having all atom types converted to carbons and all bond types converted to single bonds.^82^ Exploration of new chemical scaffolds is very import in drug discovery and development, *inter alia* with regard to the patentability. In our study, we compute the Bemis-Murcko scaffolds and cyclic skeletons of the fragments and all reference molecules using RDKit,^77^ and analyze, in case which of our fragments represent scaffolds absent from the reference datasets.

In contrast to this stringent scaffold definition, the term ‘chemotype’ is only loosely defined.^83^ We will use this term to describe molecules sharing similar structural features. Molecules of the same chemotype may, but not necessarily share the same scaffold. We additionally inspect the three top-scored and thus most similar reference molecules (irrespective from which of the reference datasets) per fragment. To this end, we computed the pairwise Tanimoto similarity for each fragment and reference molecule based on their Morgan fingerprints (radius = 2, 4096 bits) using RDKit.^77,84^

#### 5.4.3 Molecular Descriptors

Molecular Descriptors aim to express various characteristics of compounds, such as physicochemical properties, shape, topology, and molecular complexity. In the following, the descriptors used in our analyses will be introduced.

All but one of the descriptors were calculated from the molecules’ 2D structure. One additional descriptor was computed from the molecules’ 3D structure, which was either generated *in silico* or obtained from crystallographic data representing the protein-bound state. This two-legged approach was chosen to facilitate comparisons with reference datasets lacking 3D structural information, such as those derived from ChEMBL, while also leveraging available experimental 3D data where possible.

For our fragment hits as well as for the PDB reference dataset for which 3D structural information was available, we used the protonation state(s) and tautomeric form(s) of the protein-bound ligand as assigned by PROTOSS (*vide infra*). In cases where multiple ligand copies have been resolved (in any crystal structure), possibly even representing different protomers/tautomers, the descriptor values were averaged per ligand to yield a total of 160 and 36 descriptor values for the PDB reference dataset and our fragments, respectively.

##### Crystallographic Protein-Bound Conformation

We first identified the missing hydrogen atoms in the protein-ligand complex using PROTOSS,^85^ filtered the ligands based on our definition, as well as for ligand copies for which all atoms have been resolved only. For eight ligands, no fully resolved copy was available in any complex structure, namely fragments F009 and F012, and PDB ligands 46P, 7W8, 9NT, AO8, BVZ and GGB. Instead, we selected the copy with the highest number of resolved non-hydrogen atoms for those. After filtering, a total of 270 ligand instances remained in the PDB dataset and 57 fragment copies, which were subsequently subjected to descriptor calculation. The percentage of ligands for which more than one copy has been considered amounted to 39% for the fragments and 27% for the PDB ligands, respectively.

##### Computational Conformer Generation

Computational conformer generation relied on RDKit’s EmbedMolecule functionality starting from random coordinates^86^ with a fixed seed to ensure reproducibility. All other parameter settings were in concordance with the pre-configured parameter object rdkit.AllChem.ETKDGv3(). The RDKit’s ETKDG approach combines <u>e</u>xperimental torsional-angle and ring geometry preferences with additional “basic <u>k</u>nowledge” terms and the stochastic <u>d</u>istance <u>g</u>eometry-based approach.^64,65^ It has been shown to outperform other free and commercial conformer generators in producing a conformation closely resembling the bioactive conformation, typically with a root-mean-square deviation (RMSD) of less than 2 Å.^66,87^ This way, even if the generated ligand conformation may not equal the lowest energy conformation or the most stable in solution, it will closely match the conformation found in protein–ligand complexes.^66^

##### Comparison of Conformations

To compare the *in silico*-generated conformers and the crystallographic ligands, the non-hydrogen atom root-mean-square deviation (RMSD) was computed. For this purpose, an abstracted representation of the 3D molecules was used in which all non-hydrogen atoms were converted to carbon and all bond types to single bonds. This approach focuses on molecular shape rather than chemical detail, thereby avoiding mismatches.

##### Introducton to and Calculation of Descriptors

The Natural Product Likeness (NP likeness) score quantifies the overall similarity of a molecule to the structural space covered by NPs.^88^ Broadly speaking, it is calculated by comparing the structural features of a compound against the ones of a database of known NPs and normalizing for molecular size. Typically, the NP likeness score ranges from –5 to +5, with higher values indicating a greater resemblance of the molecule to NPs. While Ertl *et al.*^88^ acknowledge that NPs also differ from synthetic molecules in their global physicochemical properties, they emphasize that the major differences between these two classes lie in their structural characteristics, which is why only these features are considered in their NP likeness score. The descriptor was demonstrated to efficiently separate NPs from synthetic molecules.^88^

The fraction of sp^3^-hybridized carbon atoms (Fsp^3^) and the fraction of stereogenic carbons (FC_stereo_) are widely employed, easily interpretable descriptors for characterizing saturation and spatial complexity of molecules. These metrics are defined as the number of sp^3^-hybridized carbons or stereogenic/chiral carbons, respectively, divided by the total number of all carbons in the molecule.^7^ Lovering *et al.* demonstrated in their seminal *Escape from Flatland* publications that both measures correlate with success in high-throughput screening campaigns and overall clinical success.^7,8^ Moreover, Fsp^3^ is inversely correlated with compound promiscuity and cytochrome P450 inhibition, both associated with the risk for toxicity.^8^ When evaluating molecular 3D character using Fsp^3^, Kombo *et al.*(2013)^63^ suggested a cut-off value of *→* 0.42. In other words, at least 42% of the carbon atoms should be sp^3^-hybridized. The main limitation of both, Fsp^3^ and fraction of stereogenic carbons (FC_stereo_), is that they disregard non-carbon atoms, which may constitute a considerable portion of the molecule and contribute to its 3D structure. Consequently, these descriptors do not fully capture molecular topology and do not align with the chemical intuition of complexity. For instance, they may yield the lowest possible score of 0 even for molecules with considerable topological complexity, such as triphenylphosphine, or the highest score of 1 for relatively simple structures such as an aliphatic hydrocarbon chain.

The spatial score (SPS), introduced by Krzyzanowski *et al.*, combines the benefits of both Fsp^3^ and FC_stereo_, but also overcomes shortcomings caused by their simplistic nature.^89^ It is based on four parameters, calculated for each non-hydrogen atom in a molecule and are then summed across the whole structure: an atom hybridization term, a stereoisomeric term (strongly related to FC_stereo_), a non-aromatic ring term and the number of directly bonded non-hydrogen atoms.^89^ As the SPS depends on molecular size, the authors suggested normalizing the score by dividing by the total number of non-hydrogen atoms. The resulting nSPS was reported to correlate with biologically relevant properties, such as selectivity and potency.^89^

The deviation from the plane of best fit (PBF) descriptor was the only descriptor computed from the molecules’ 3D structures. Notably, as the conformers generated may differ amongst different software solutions, values cannot be directly compared between publications. The PBF descriptor measures the sum of the distances of all non-hydrogen atoms from the plane of best fit.^90^ In general, a higher PBF value corresponds to a higher degree of three-dimensionality. While its theoretical range extends from zero to infinity, in practice, the PBF value tends to be *<* 2 for small drug-like molecules even if they are rich in 3D-character. Two examples of particularly globular or planar molecules are the natural products and approved drugs adamantine and salicylic acid, for which PBF values of 1.02 and 0.13 are obtained. However, to account for its size-dependency, the PBF value should be either only used to compare similarly sized compounds or should be normalized, *e. g.* divided by the number of non-hydrogen atoms, which was also applied in this study. The two example molecules feature nPBF values of 0.093 and 0.013, respectively.

While Fsp^3^, FC_stereo_ and normalized spatial score (nSPS) descriptors have originally been introduced as measures of saturation and the molecular (spatial) complexity,^7,89^ nPBF expresses the molecular shape/three-dimensionality.^90,91^ Notably, it has been shown that Fsp^3^ and PBF only show poor correlation.^90,92^ Firth *et al.*proposed that this might be explained by the fact that while a high Fsp^3^ value indicates a high ratio of sp^3^-hybridized carbons to all carbons, a high Fsp^3^ score does not characterize whether these sp^3^-hybridized carbons are connected to extended vectors out of the plane of the dominant ring system; *i.e.*, contribute to significant 3D shape.

##### Descriptor Analysis

On the one hand, we analyzed the descriptor values of all ligands, on the other hand, we made use of the binding site annotation determined earlier (*cf.* section 5.3) and examined the subdataset of ATP site ligands.

Radar plots facilitate the comparison of the median values across the multitude of descriptors at first glance. Moreover, a combination of swarm plots and rain cloud plots is used to visualize the descriptor value distributions for each dataset. The latter combine a half violin plot (kernel density estimate, ‘cloud’) with jittered raw data (‘rain’) and the corresponding box plot (’umbrella’). To assess whether the descriptor values differed significantly across datasets, first, the non-parametric Kruskal–Wallis test was employed.^93^ As it indicated significant differences, Dunn’s post-hoc test^94^ was used to perform pairwise comparisons between the fragments and the reference datasets for each descriptor. Descriptor differences were considered statistically significant if the p-value was below 0.05. The tests were implemented using scipy.stats.kruskal and scikit_posthocs.posthoc_dunn functions.

## Ancillary Information

### Associated Content

#### Data Availability

All data and code underlying this study are openly available from GitHub (NPFrag2Kinase) and the PDB. The reference datasets were compiled from sources in the public domain, namely COCONUT, PDB, ChEMBL and KLIFS databases, as described in the methods section, and are also stored on GitHub.

#### Supporting Information

- Additional figures and tables, e. g. descriptor value distributions, etc. (PDF)
- X-ray and crystallographic data (XLSX)
- Fragment SMILES (CSV)
- Fragment copy information (SDF), incl. binding site annotations, *in silico*-generated conformation and computed descriptor values.

#### Accession Codes

The authors will release the atomic coordinates of all 36 disclosed PDB structures upon article publication: F001 (9RDU), F005 (9RDV), F009 (9RDW), F012 (9RDX), F024 (9RDY), F030 (9RDZ), F032 (9RE0), F055 (9RE1), F058 (9RE2), F070 (9RE3), F073 (9RE4), F074 (9RE5), F102 (9RE6), F134 (9RE7), F138 (9RE8), F145 (9RE9), F168 (9REA), F184 (9REB), F186 (9REC), F188 (9RED), F189 (9REE), F203 (9REF), F225 (9REG), F236 (9REH), F248 (9REI), F264 (9REJ), F274 (9REK), F283 (9REL), F294 (9REM), F296 (9REN), F299 (9REO), F304 (9REP), F310 (9REQ), F312 (9RER), F313 (9RES), F322 (9RET).

## Acknowledgements

The authors thank the Helmholtz-Zentrum Berlin für Materialien und Energie for the allocation of synchrotron radiation beamtime. In particular, they thank Manfred Weiss. The authors acknowledge the use of artificial intelligence tools, namely ChatGPT, BlackboxAI, and DeepL, which supported language refinement and coding assistance. These tools were not used for the *de novo* generation of text or content conceptualization.

## Abbreviations Used

3D: three-dimensional
ATP: adenine triphosphate
COCONUT: COlleCtion of Open NatUral producTs
EDIA: electron density score for individual atoms
EDIA_m_: electron density score for multiple atoms
FBDD: fragment-based drug discovery
FC_stereo_: fraction of stereogenic carbons
Fsp^3^: fraction of sp^3^-hybridized carbon atoms
NP: natural product
nPBF: normalized deviation from the plane of best fit
nSPS: normalized spatial score
PDB: Protein Data Bank
PKA: protein kinase A
PKI: protein kinase inhibitor
RMSD: root-mean-square deviation

