## Supplementary material for "Natural Product-Like Fragments Unlock Novel Chemotypes for a Kinase Target – Exploring Options beyond the Flatland": SI

### Supporting Information

#### Contents

S 1 Binding Modes of Fragment Hits . . . . . . . . . . . . . . . . . . . . . . . . . . . . . . SI 2
S 2 Natural Products Comprising The Fragments As Substructures . . . . . . . . . . . . . . SI 4
S 3 Allosteric Pocket Occupation . . . . . . . . . . . . . . . . . . . . . . . . . . . . . . . SI 5
S 3.1 F189 Binds Specifically at Peripheral Site E . . . . . . . . . . . . . . . . . . . . . . SI 6
S 4 Chemical Space Analysis . . . . . . . . . . . . . . . . . . . . . . . . . . . . . . . . . SI 8
S 4.1 Bemis-Murcko Scaffolds . . . . . . . . . . . . . . . . . . . . . . . . . . . . . . . . SI 8
S 4.2 Chemotypes . . . . . . . . . . . . . . . . . . . . . . . . . . . . . . . . . . . . . . SI 10
S 4.3 Molecular Descriptors . . . . . . . . . . . . . . . . . . . . . . . . . . . . . . . . . SI 17

### S 1 Binding Modes of Fragment Hits

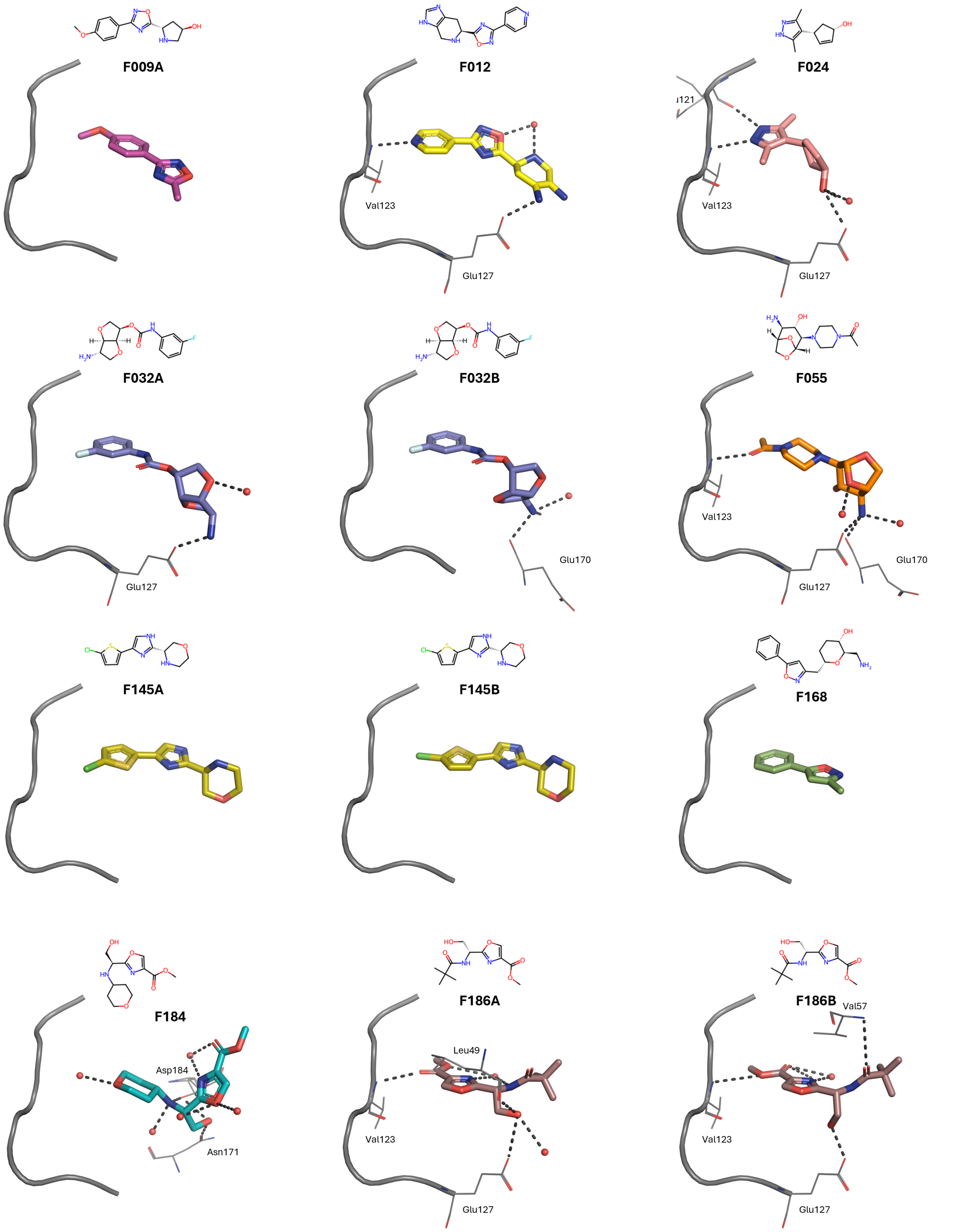


**Figure S1:** Binding modes of fragments within PKA's ATP pocket. If multiple conformations of the fragments were resolved, we show each individually (A/B). Fragments are shown as sticks, amino acid residues involved in polar interactions as lines, waters involved in polar interactions as red spheres, and polar interactions as dashed lines. Additionally, the kinase hinge and linker (residues 120 – 127) are shown in cartoon representation. For clarity, only amino acid residues involved in polar interactions are labeled.

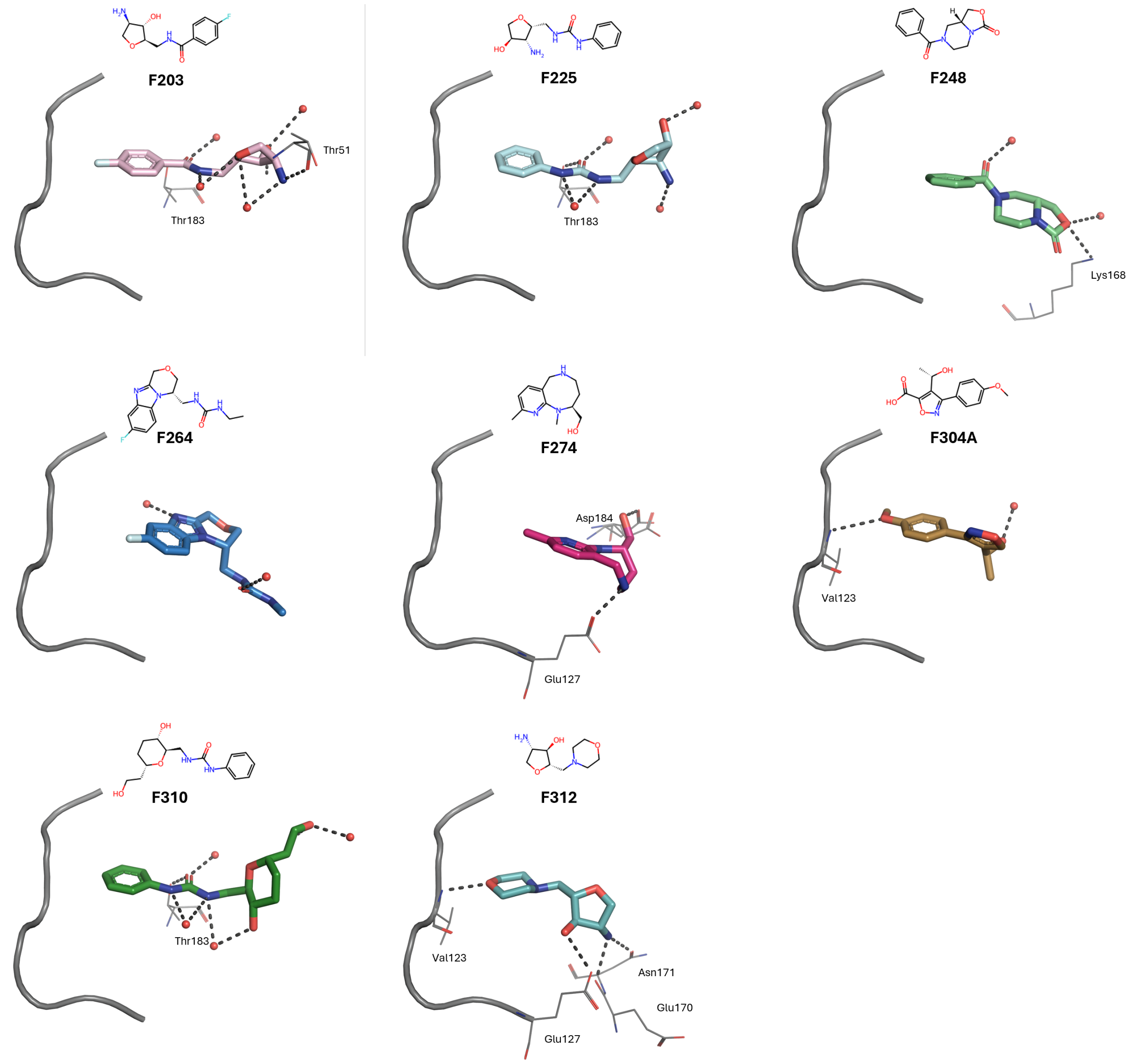


**Fig. S1**: Continued.

#### S 2 Natural Products Comprising The Fragments As Substructures

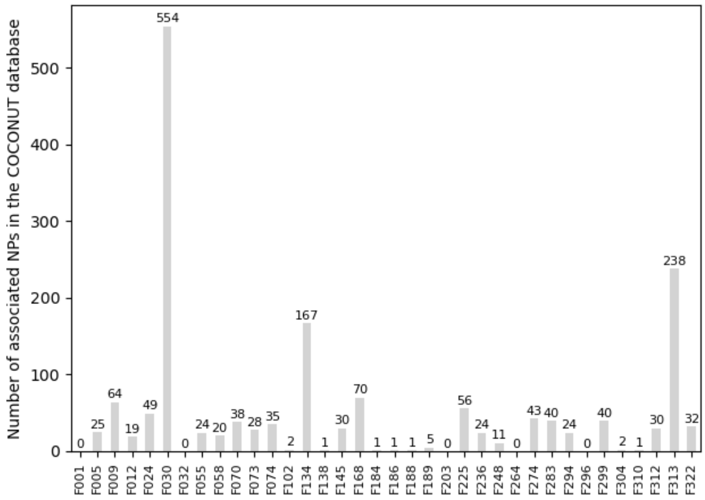


**Figure S2:** Number of Natural Products (NPs) from the COCONUT database, comprising our fragment hits as a substructure. Unsurprisingly, all the non-fluorinated fragments,*i.e.*, all fragments except F001, F032, F203, F264 and F296, were found to possess an entry in the COCONUT database themselves, as the latter sources molecules from the subset of NPs in the ZINC15 database, among others, which in turn integrates the AnalytiCon Discovery libraries.

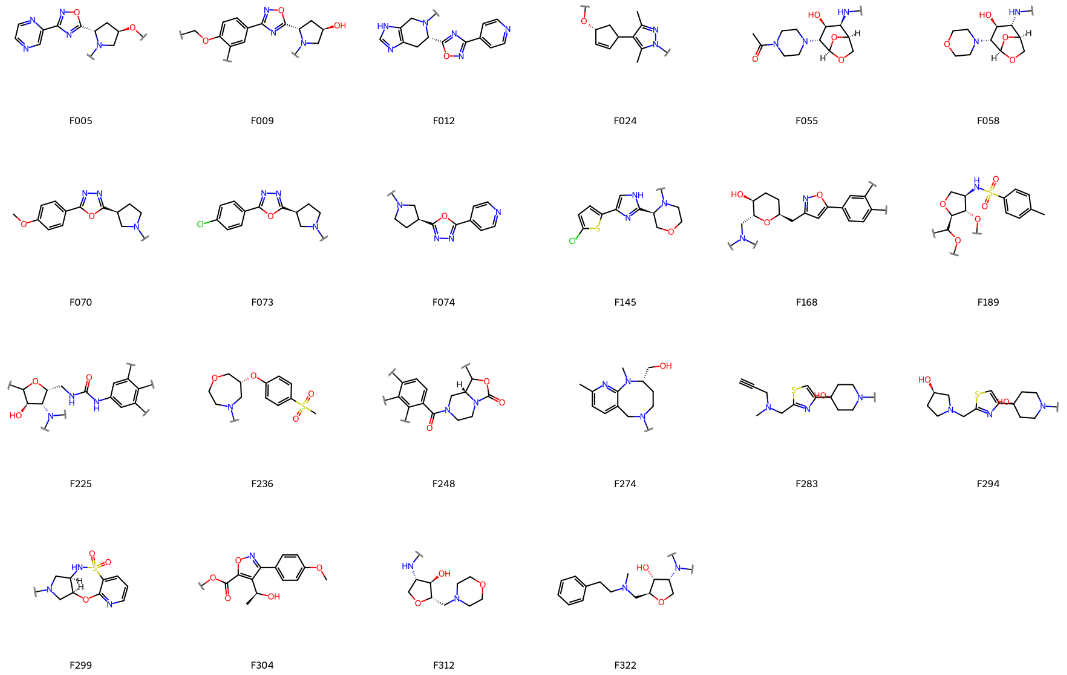


**Figure S3:** Growth vectors in the fragments, as deduced from the derivatization patterns of the associated natural products, indicated by a squiggled line perpendicular to a bond.

#### S 3 Allosteric Pocket Occupation

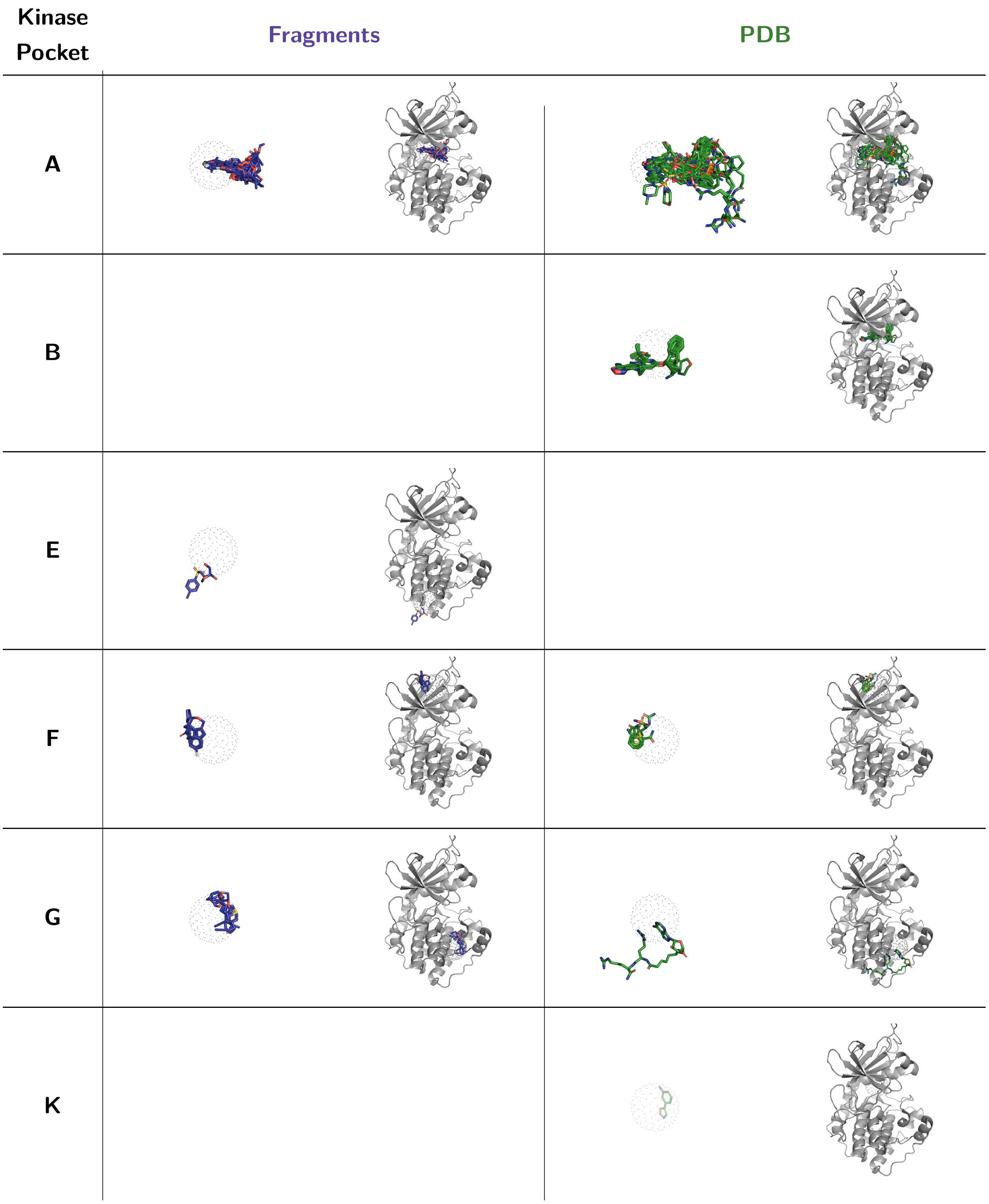


**Figure S4:** Allosteric pocket occupation by PKA ligands from the PDB and our fragment hits (stick representation with green and blue carbons, respectively). The pocket occupation of a ligand/fragment was defined as 'being located in a 5 Å radius (visualized as dots) of an added and well-considered pseudo atom' which in turn was placed in the center of mass of all ligands binding with this pocket as listed by Xerxa *et al.*.[37] Thus, the position of ligands occupying the same pocket can be slightly shifted. For reference, the PKA structure (PDB-ID: 3FJQ) is shown in ribbon representation.

| Kinase Pocket | Fragments | PDB |
| --- | --- | --- |
| Other | 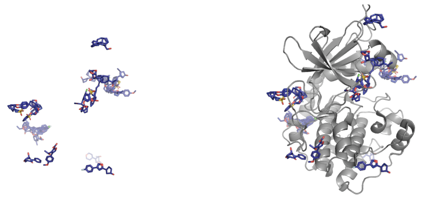 | 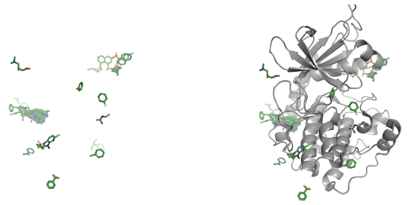 |

**Figure S4.:** Continued.

#### S 3.1 F189 Binds Specifically at Peripheral Site E

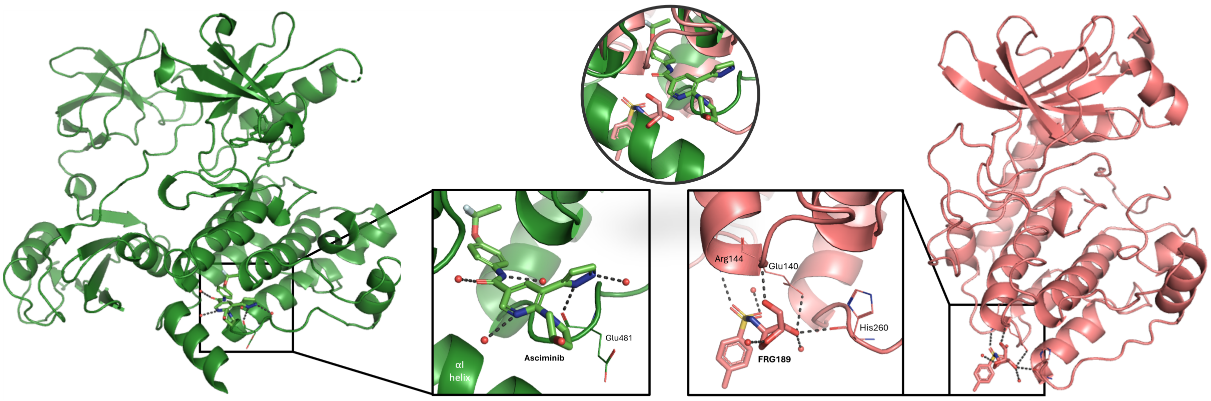


**Figure S6:** Comparison of binding modes for asciminib and F189. Ligands are shown as sticks, both kinases are shown in cartoon representation, hydrogen bonds are visualized as dashed black lines. Moreover, in the insets, amino acid residues involved in polar interactions are depicted as lines and water molecules as red spheres. For clarity, we show both complex structures and binding modes independently, but in the same orientation. **Left:** Asciminib bound to BCR-ABL1 (PDB-ID: 5MO4). The C-terminal αI-helix of tyrosine kinases such as ABL1 exhibits significant conformational flexibility and borders the allosteric pocket E in its kinked conformation,[57] which is not the case for PKA. **Right:** F189 bound to PKA. **Middle:** In addition, we show the superimposition of asciminib bound to BCR-ABL1 and F189 bound to PKA in the circular inset. Remember, that the pocket occupation of a ligand/fragment was defined as 'being located in a 5 Å radius of an added and well-considered pseudo atom' which in turn was placed in the center of mass of all ligands binding with this pocket as listed by Xerxa *et al.*[37] (*cf.* 5.3 Analysis of Ligand-Bound Protein Kinase A Crystal Structures). Thus, the position of ligands occupying the same pocket can be slightly shifted.

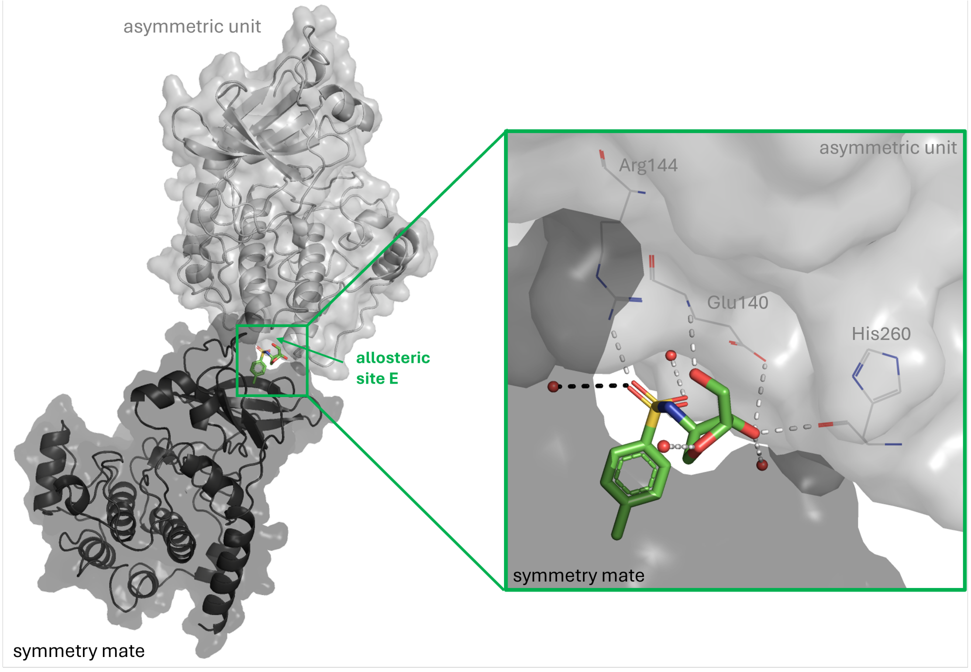


**Figure S7:** Hydrogen bonding interactions of F189 (stick representation with green carbons) within the asymmetric unit and with the neighboring symmetry mate in the crystal lattice (surface/cartoon representation, light gray and black, respectively). Hydrogen bonds are visualized as dashed white and black lines, respectively; the protein residues and water molecules involved in them are displayed in line representation or as spheres.

### S 4 Chemical Space Analysis

#### S 4.1 Bemis-Murcko Scaffolds

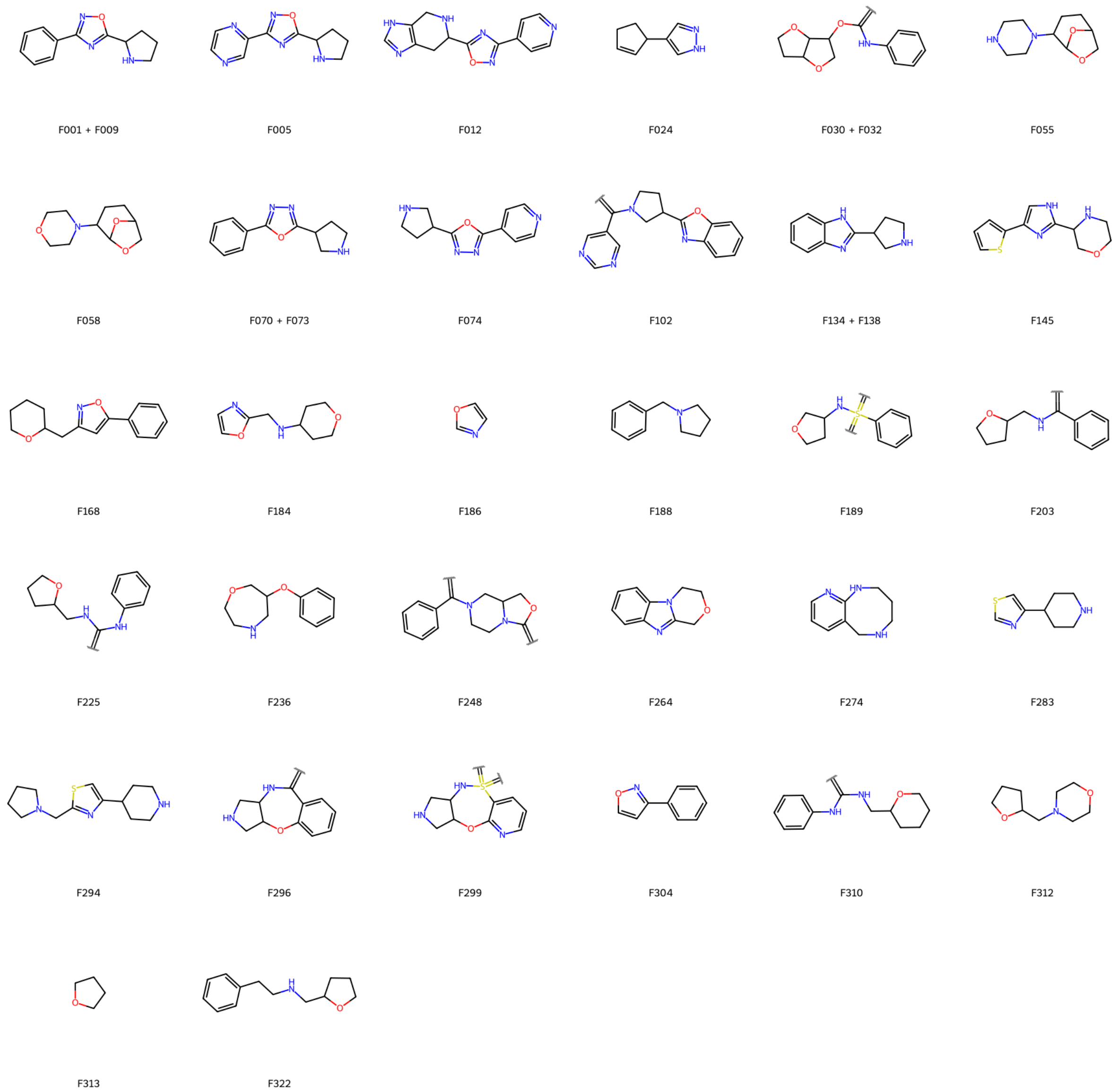


**Figure S8:** Bemis-Murcko scaffolds of our fragments. None of them is found in the reference datasets.

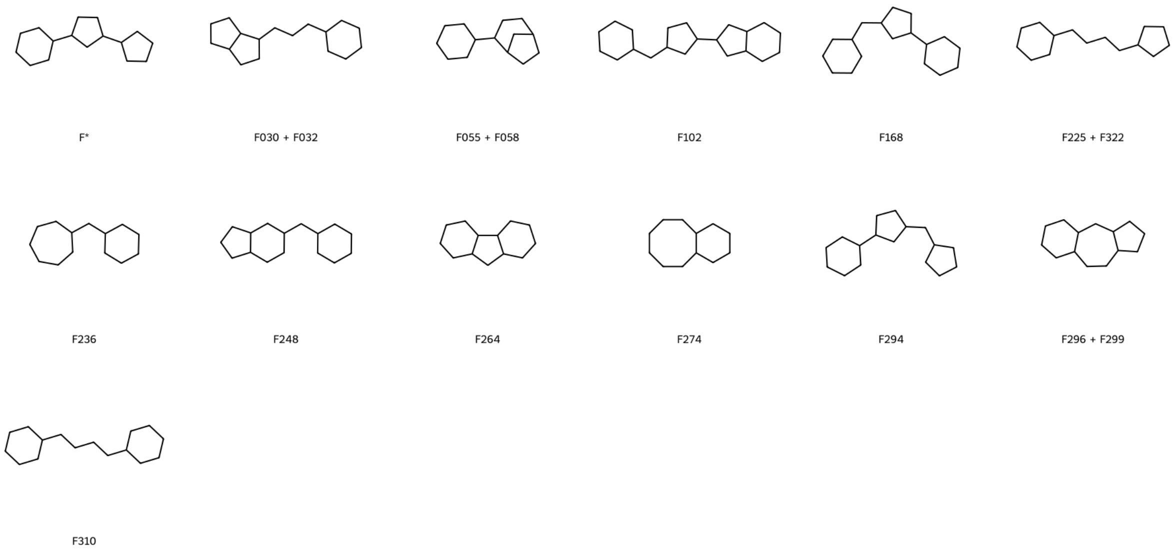


**Figure S9:** Cyclic skeletons of our fragments, not found in any of the PKA-specific reference datasets. F * = F001 + F005 + F009 + F070 + F073 + F074 + F145.

#### S 4.2 Chemotypes

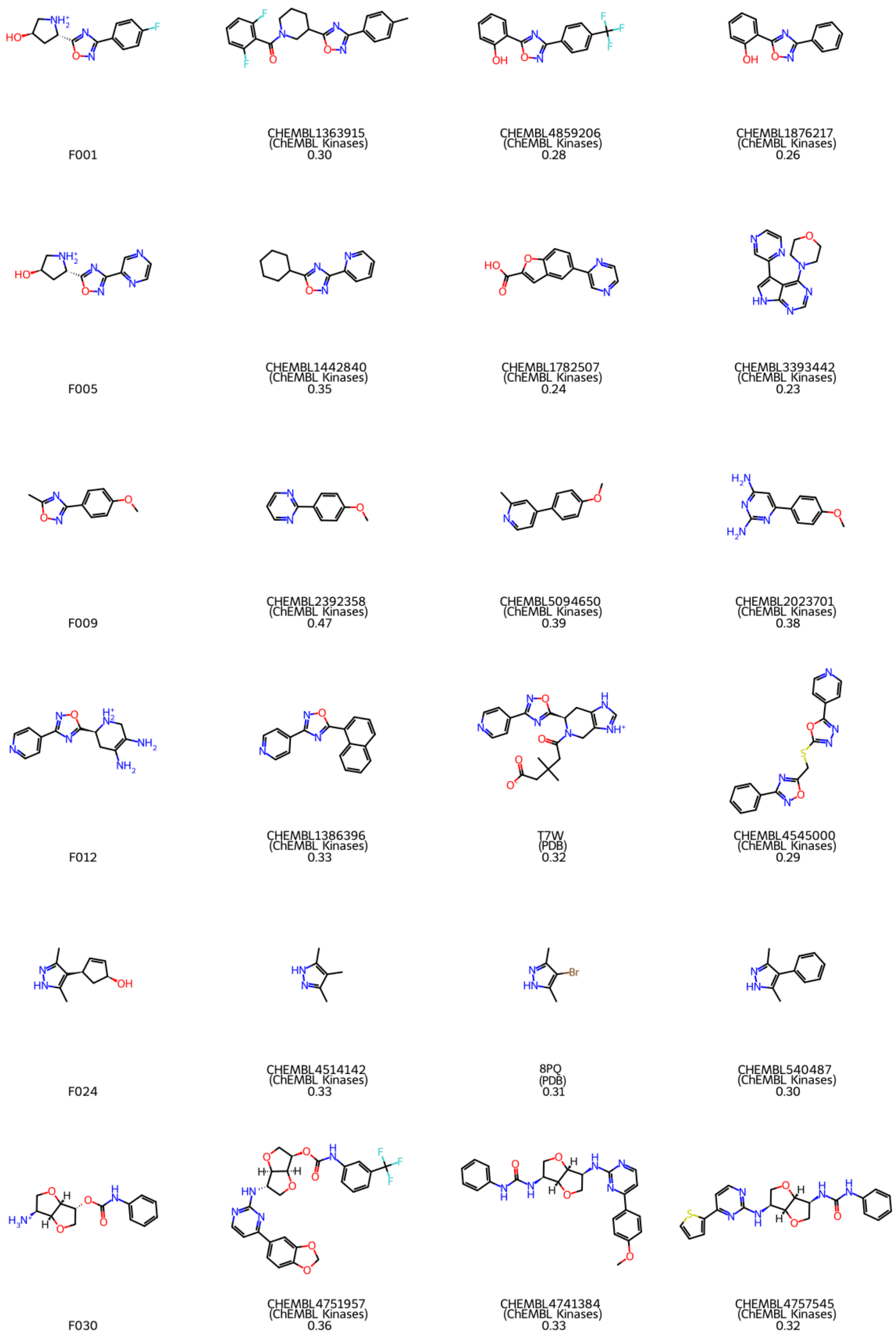


**Figure S10:** In each row, the fragment structure is shown on the left, along with the three most similar reference molecules from any of the reference datasets to the right. Underneath each reference molecule, the molecule identifier, the reference data set from which it originates and the respective Tanimoto coefficient are printed.

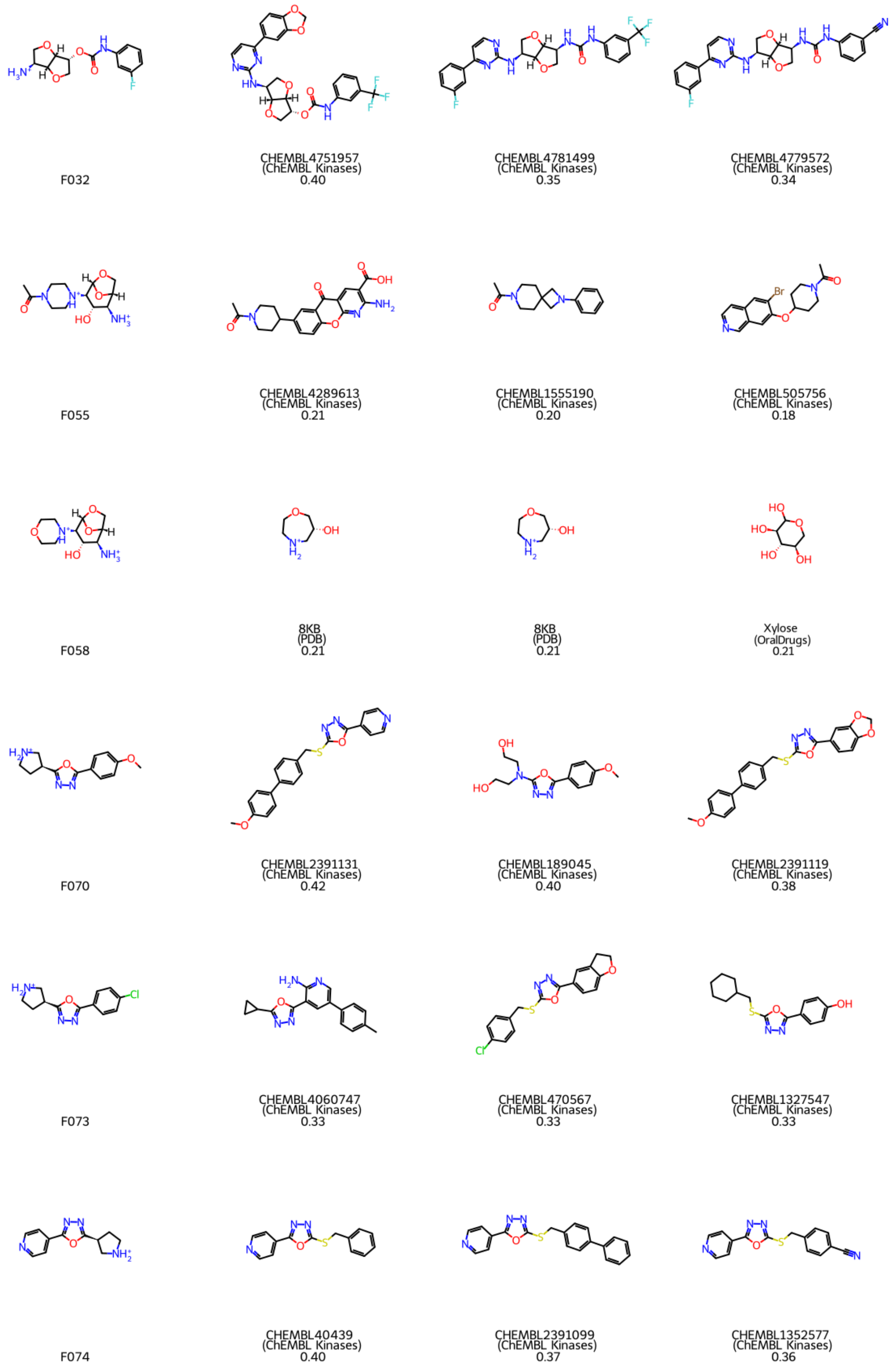


**Figure S10:** Continued.

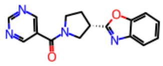

F102

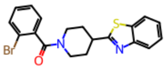

CHEMBL1387577
(ChEMBL Kinases)
0.33

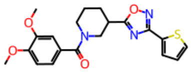

CHEMBL1459223
(ChEMBL Kinases)
0.32

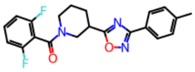

CHEMBL1363915
(ChEMBL Kinases)
0.31

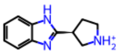

F134

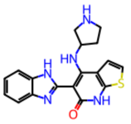

CHEMBL3330134
(ChEMBL Kinases)
0.39

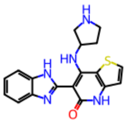

CHEMBL3330237
(ChEMBL Kinases)
0.39

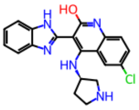

CHEMBL210035
(ChEMBL Kinases)
0.37

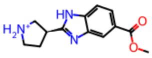

F138

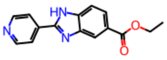

CHEMBL4638916
(ChEMBL Kinases)
0.39

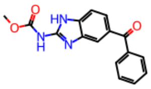

Mebendazole
(OralDrugs)
0.37

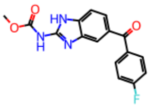

CHEMBL1454946
(ChEMBL Kinases)
0.37

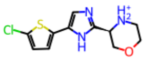

F145

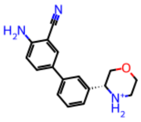

7QI
(PDB)
0.27

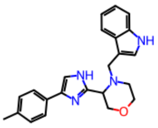

CHEMBL2297166
(ChEMBL Kinases)
0.27

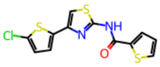

CHEMBL1308898
(ChEMBL Kinases)
0.27

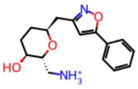

F168

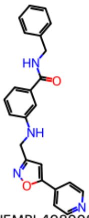

CHEMBL4089087
(ChEMBL Kinases)
0.28

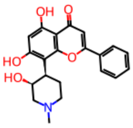

CHEMBL3898028
(ChEMBL Kinases)
0.24

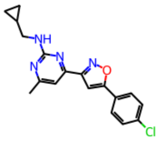

CHEMBL1377267
(ChEMBL Kinases)
0.24

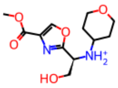

F184

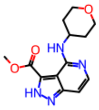

CHEMBL3651815
(ChEMBL Kinases)
0.28

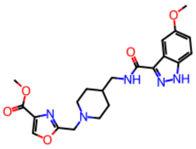

CHEMBL3735500
(ChEMBL Kinases)
0.28

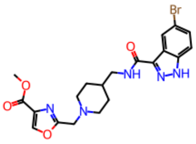

CHEMBL3736476
(ChEMBL Kinases)
0.28

**Figure S10:** Continued.

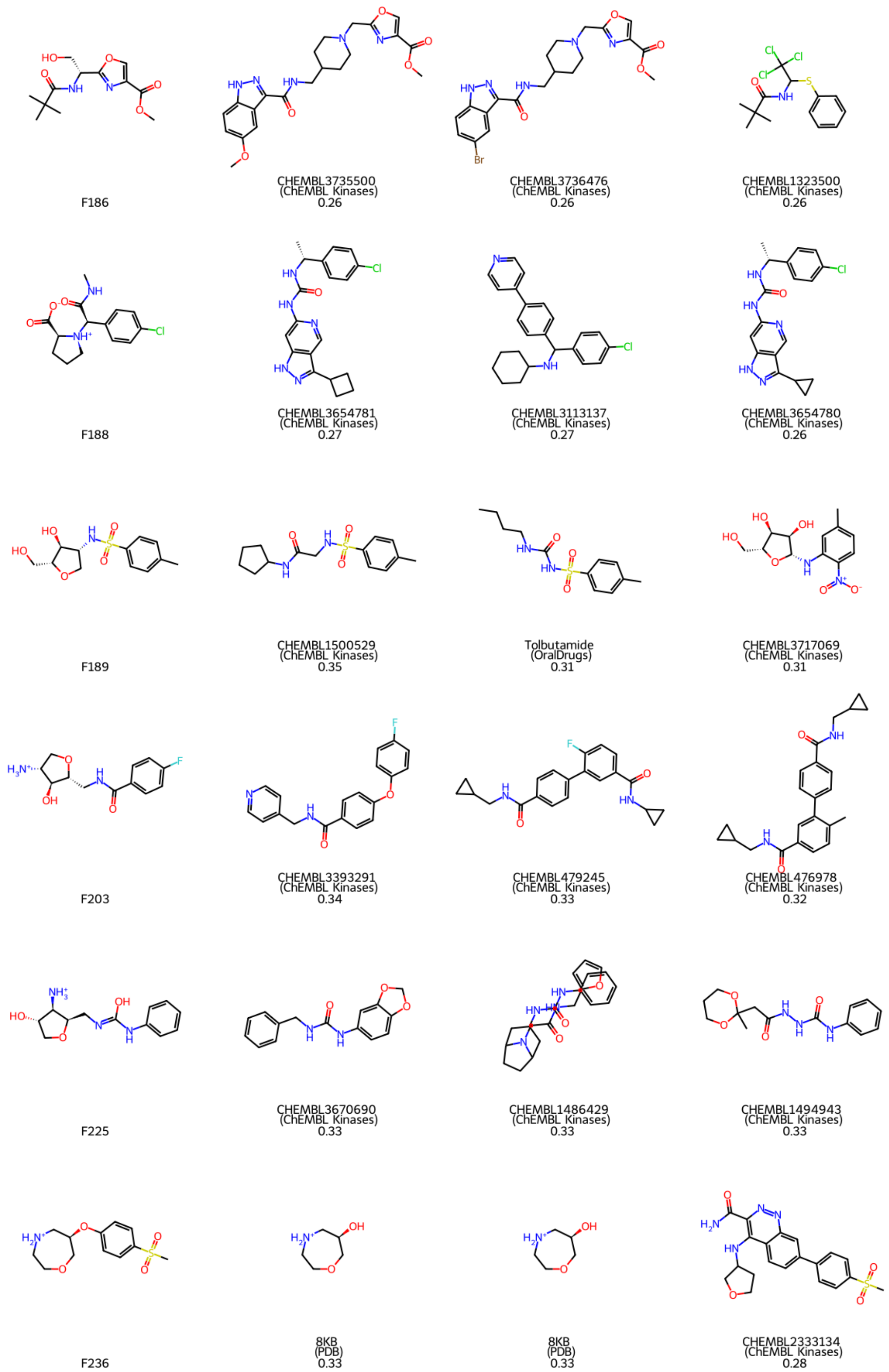


**Figure S10:** Continued.

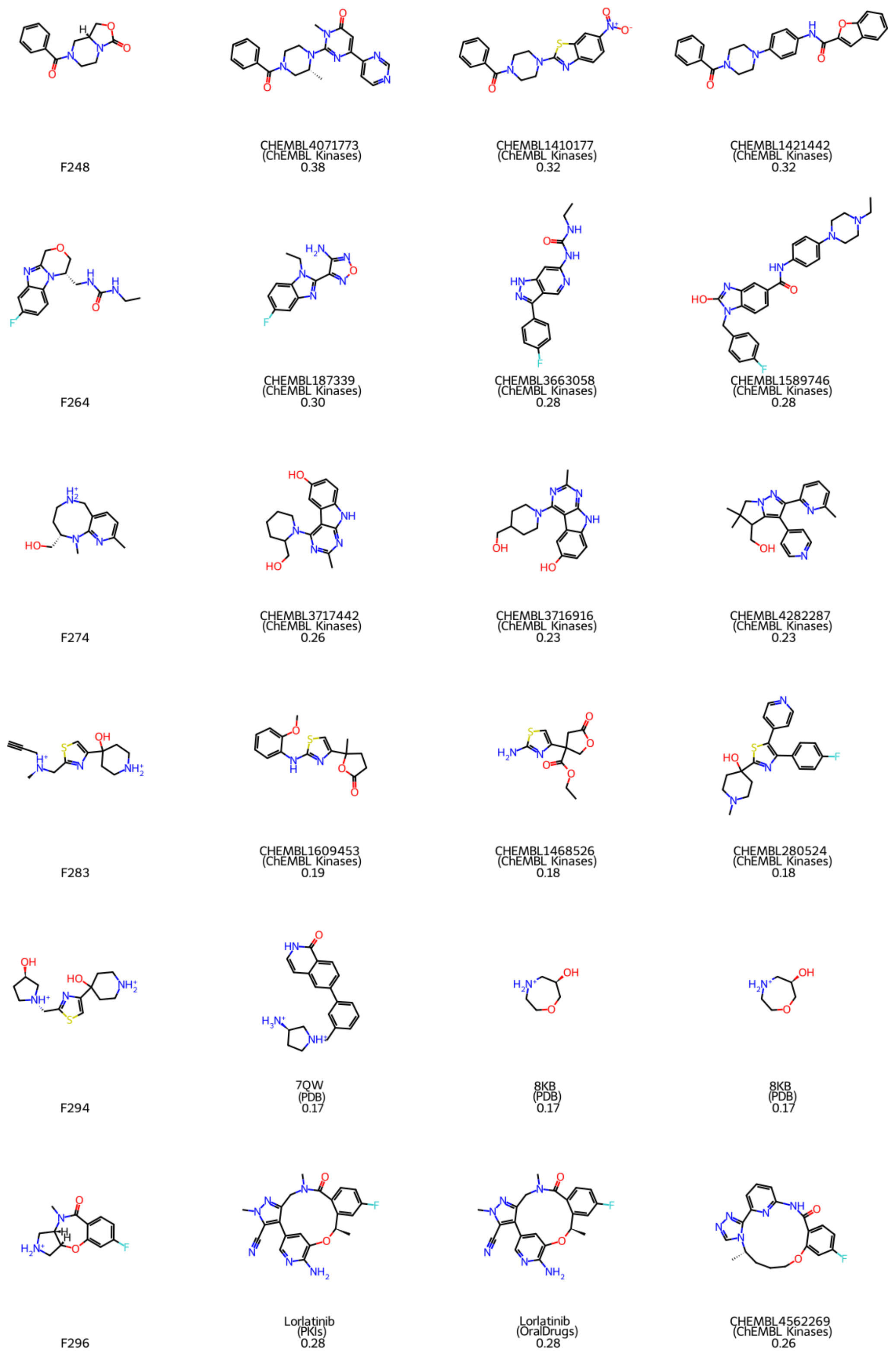


**Figure S10:** Continued.

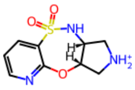

F299

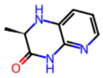

CHEMBL3984982
(ChEMBL Kinases)
0.22

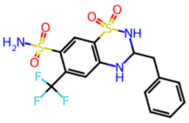

Bendroflumethiazide
(OralDrugs)
0.21

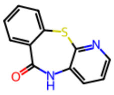

CHEMBL1613269
(ChEMBL Kinases)
0.21

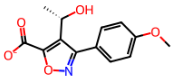

F304

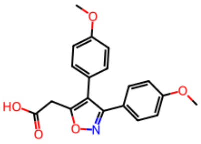

Mofezolac
(OralDrugs)
0.46

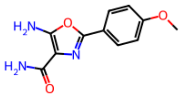

CHEMBL208516
(ChEMBL Kinases)
0.35

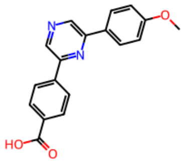

CHEMBL3359882
(ChEMBL Kinases)
0.33

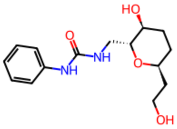

F310

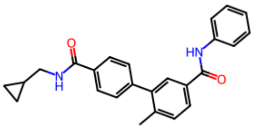

CHEMBL516264
(ChEMBL Kinases)
0.31

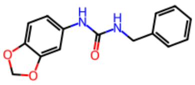

CHEMBL3670690
(ChEMBL Kinases)
0.31

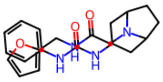

CHEMBL1486429
(ChEMBL Kinases)
0.31

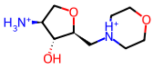

F312

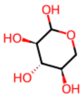

Xylose
(OralDrugs)
0.22

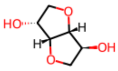

Isosorbide
(OralDrugs)
0.20

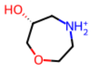

8KB
(PDB)
0.20

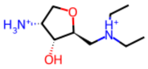

F313

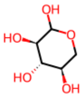

Xylose
(OralDrugs)
0.24

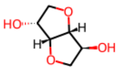

Isosorbide
(OralDrugs)
0.21

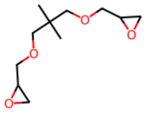

CHEMBL1587512
(ChEMBL Kinases)
0.20

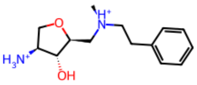

F322

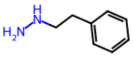

Phenelzine
(OralDrugs)
0.24

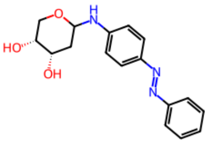

CHEMBL1445664
(ChEMBL Kinases)
0.24

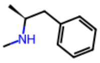

Methamphetamine
(OralDrugs)
0.23

**Figure S10:** Continued.

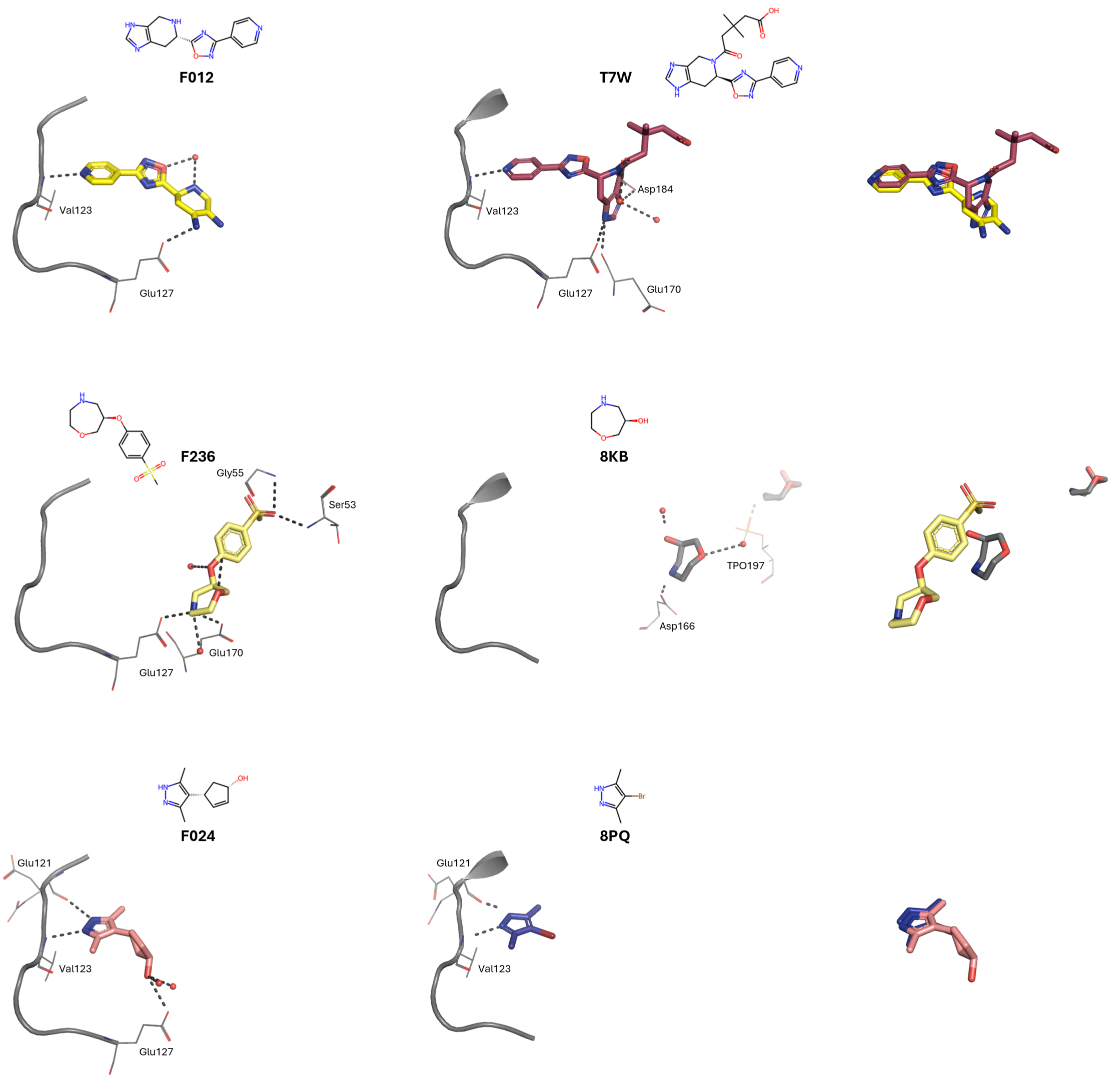


**Figure S11:** Comparison of Binding Modes for the fragments F012, F236 and F024 and their most similar PKA ligands from the PDB, namely 7TW (PDB-ID: 7BB0, Tanimoto similarity of 0.46 with F012), 8KB (PDB-ID: 5N3G, Tanimoto similarity of 0.33 with F236), 8PQ (PDB-ID: 5N7U, Tanimoto similarity of 0.31 with F024). Fragments are shown as sticks, amino acid residues involved in polar interactions as lines, waters involved in polar interactions as red spheres, and polar interactions as dashed lines. Additionally, the kinase hinge and linker (residues 120 – 127) are shown in cartoon representation. For clarity, only amino acid residues involved in polar interactions are labeled. On the right, we show the superimposition of the related ligands. T7W comprises F012 as a substructure, but differs in stereochemistry. Despite this difference, both molecules bind to the same site of the ATP pocket and form a hydrogen bond between the pyridine-N and the backbone amide-NH of the hinge residue Val123, as well as towards Glu127. 8KB is a substructure of F236. Their binding sites are laterally shifted, resulting in different interactions. While the secondary amine of F236 forms a salt bridge with Glu127 and Glu170, 8KB's amino functionality forms a salt bridge to Asp166. 8PQ is an analog of F024 with a bromine substituent in the 4-position of the pyrazole ring, compared to the larger cyclopent-2-en-1-ol moiety in F024. One of the two alternative conformations resolved for 8PQ (B, occupancy = 0.4) closely matches the binding mode of F024, forming hydrogen bonds via the pyrazole nitrogens to the backbone amide-NH of Val123 and the backbone carbonyl-O of Glu121. Moreover, PDB ligand 7QI (PDB-ID: 7PID, Tanimoto similarity of 0.28 with F145) appears in the list of most similar reference molecules, however, 7QI and F145 do not belong to the same chemotype, as they only a small proportion of their structure, and are therefore not compared with regard to their binding modes.

#### S 4.3 Molecular Descriptors

**Table S1:** Median values of molecular descriptors. n.d. = not determined.

| | MW | HBA | HBD | LogP | NRB | TPSA | NAR | NP-Likeness | NHA | $Fsp^3$ | $FC_{stereo}$ | nSPS | nPBF (*in silico*) | nPBF (protein-bound) |
|---|---|---|---|---|---|---|---|---|---|---|---|---|---|---|
| **Binding Site Agnostic** | | | | | | | | | | | | | | |
| Fragments | 253.8 | 4 | 2 | -0.17 | 4 | 73 | 1 | -0.46 | 18 | 0.43 | 0.12 | 23 | 0.046 | 0.041 |
| PDB | 254.3 | 3 | 2 | 1.39 | 4 | 71 | 2 | -0.92 | 19 | 0.22 | 0.00 | 12 | 0.036 | 0.033 |
| ChEMBL Kinases | 408.5 | 6 | 2 | 3.74 | 6 | 87 | 3 | -1.23 | 29 | 0.24 | 0.00 | 14 | 0.033 | n.d. |
| ChEMBL PKA | 371.4 | 5 | 2 | 3.52 | 6 | 77 | 3 | -1.12 | 27 | 0.22 | 0.00 | 15 | 0.04 | n.d. |
| Oral Drugs | 349.4 | 4 | 2 | 2.88 | 7 | 74 | 2 | -0.47 | 24 | 0.38 | 0.05 | 16 | 0.039 | n.d. |
| PKIs | 467.8 | 7 | 2 | 4.10 | 8 | 92 | 3 | -1.27 | 33 | 0.28 | 0.00 | 15 | 0.034 | n.d. |
| COCONUT NPs | 376.4 | 6 | 2 | 2.47 | 7 | 86 | 2 | -0.97 | 27 | 0.43 | 0.16 | 24 | n.d. | n.d. |
| **ATP Site Ligands** | | | | | | | | | | | | | | |
| Fragments | 250.7 | 4 | 2 | 0.40 | 4 | 72 | 2 | -0.87 | 18 | 0.42 | 0.09 | 22 | 0.043 | 0.037 |
| PDB | 273.4 | 3 | 2 | 1.44 | 4 | 72 | 2 | -0.92 | 20 | 0.22 | 0.00 | 12 | 0.036 | 0.032 |

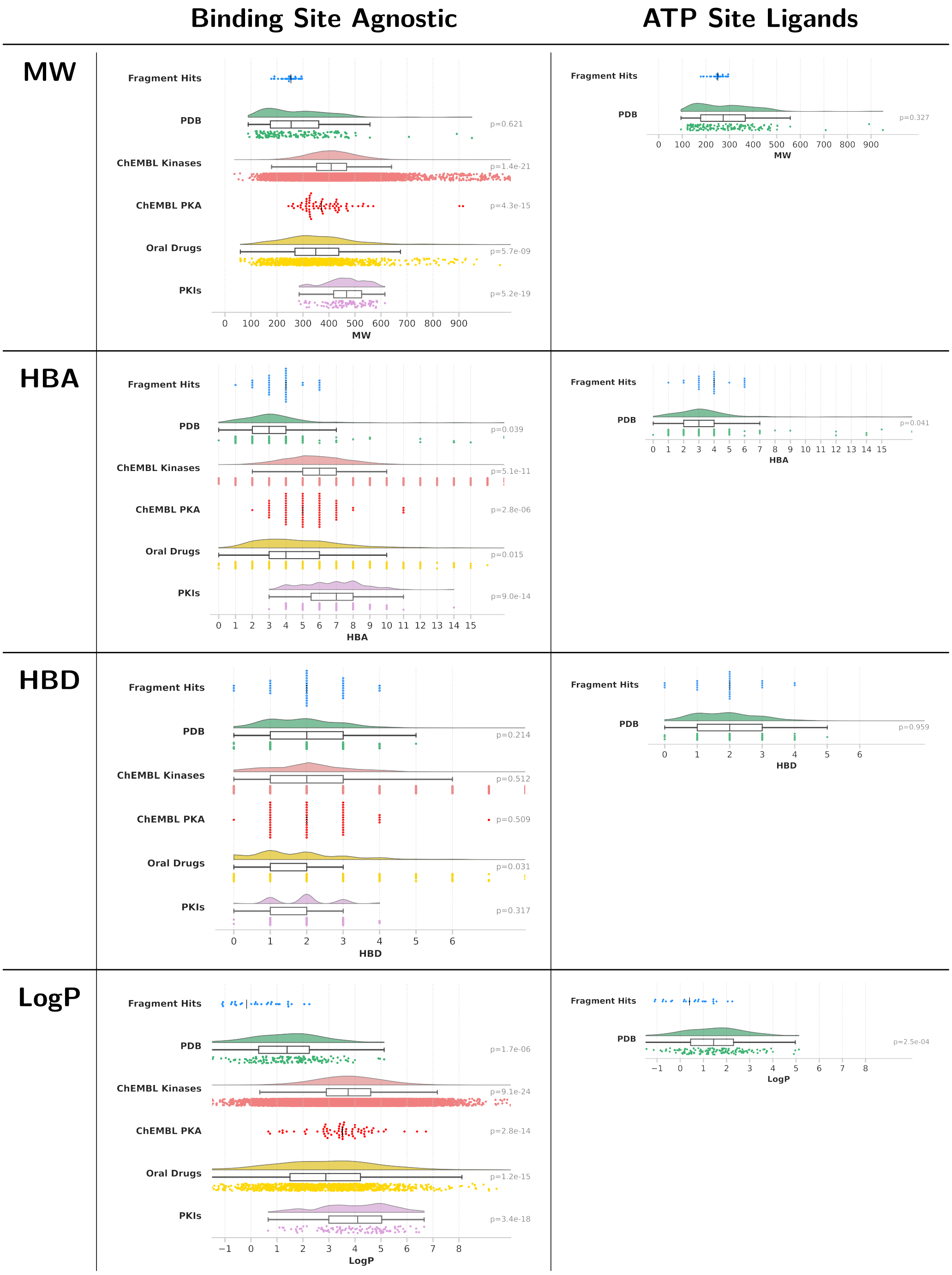


**Figure S12:** Distributions of descriptor values across the datasets, visualized as rain cloud and swarm plots, respectively, depending on the size of the dataset. In the swarm plots, the median is indicated by a vertical black line. As a guide value for the X-axis limits, the range in which 99% of all the data falls was used. For the subset of ATP-site ligands, we can only compared the datasets with structural information available, namely the fragments and the PDB reference dataset. p-values as obtained from Dunn's statistical test, comparing the fragments and the reference datasets, are shown on the right. Median values are summarized in Tab. S1. MW = Molecular weight. HBA and HBD = Number of hydrogen acceptor and donor atoms, respectively. LogP = calculated logarithm of the n-octanol/water partition coefficient.

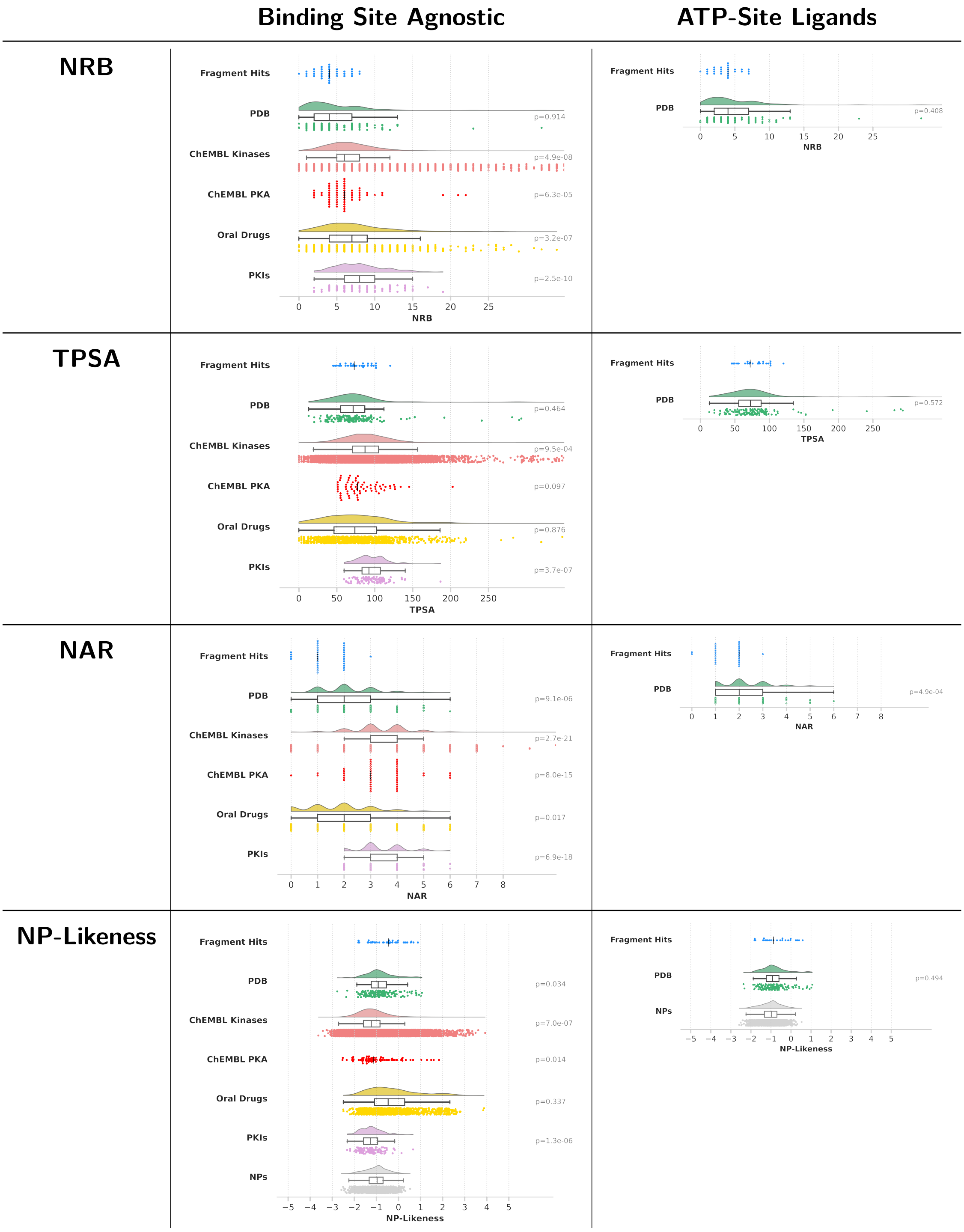


**Figure S12:** Continued. NRB = Number of rotatable bonds. TPSA = Topological polar surface area. NAR = Number of aromatic rings. NP-Likeness = Natural product likeness.

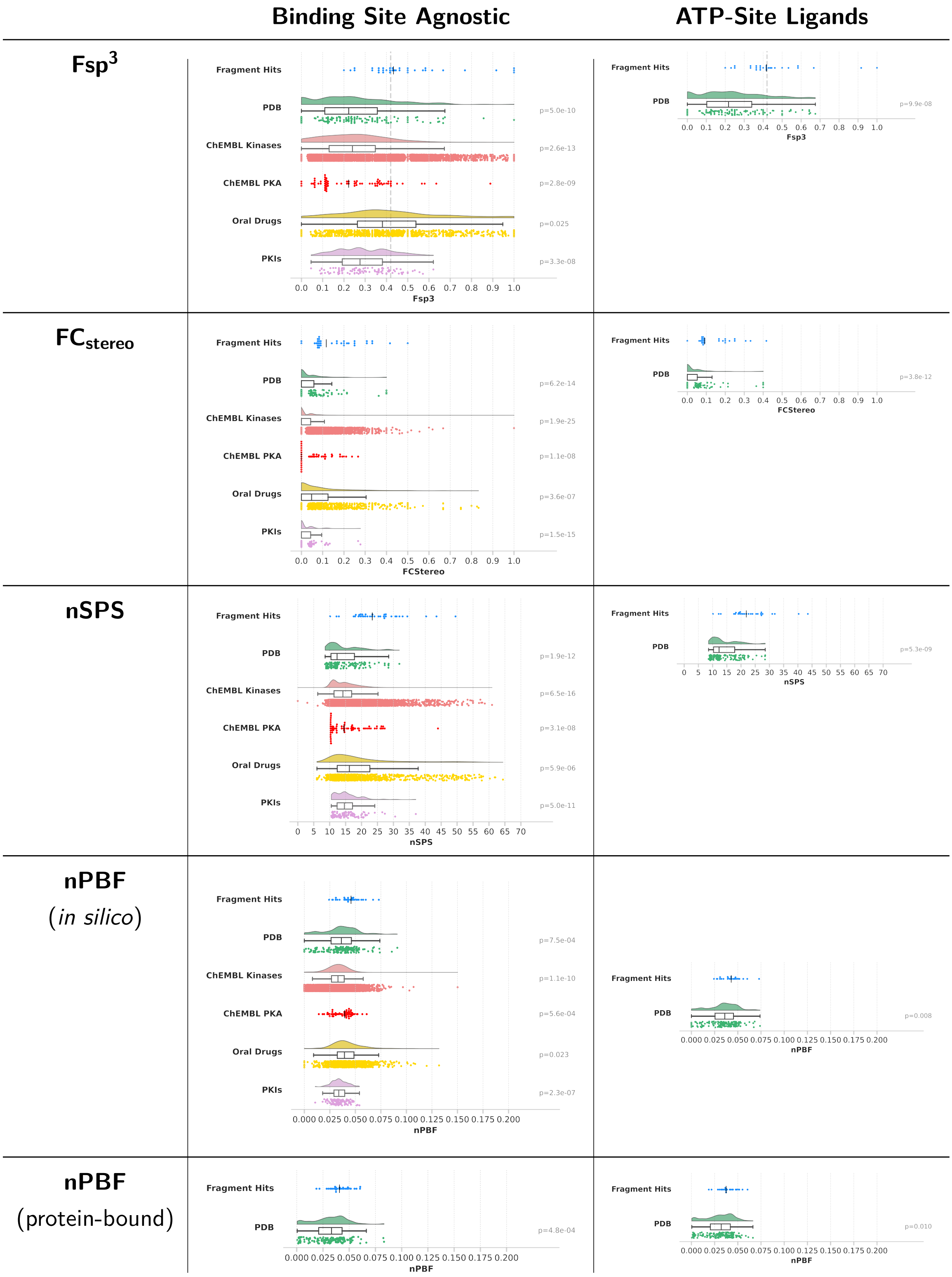


**Figure S12:** Continued. $Fsp^3$ = Fraction of $sp^3$-hybridized carbons. $FC_{stereo}$ = Fraction of stereogenic carbons. nSPS = Normalized spatial score. nPBF = Normalized deviation from the plane of best fit, calculated from *in silico*-generated or the protein-bound ligand conformations.

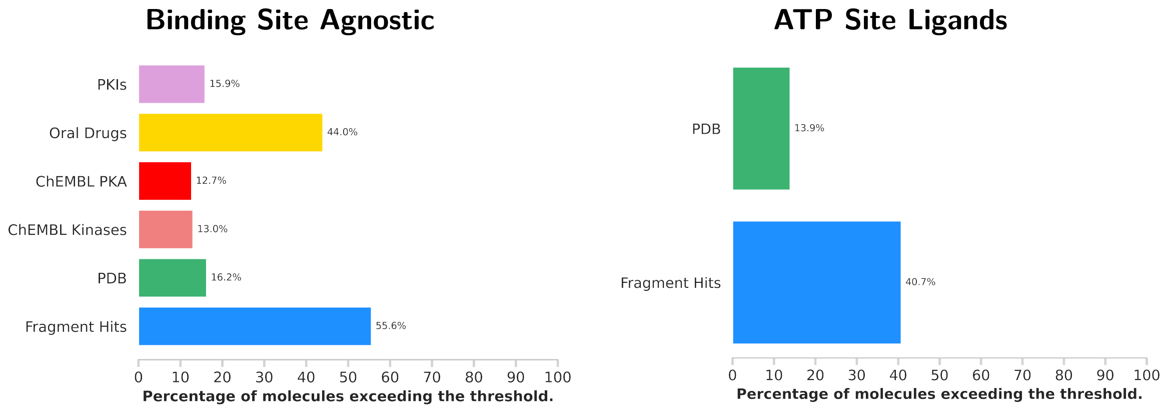


**Figure S13:** Percentage of molecules exceeding the threshold of $Fsp^3 \geq 0.42$ defined by Kombo *et al.*.[63]

**Table S2:** Molecules exhibiting the highest/lowest descriptor value per dataset. '...' indicates that more than 5 molecules possess the exact same value. n.d. = not determined. A color code similar to the one used in Fig. 1 is used: Fragments that bind only in the ATP pocket are highlighted in green; fragments that bind only at one or more peripheral site(s) are highlighted in blue; fragments binding in the ATP pocket and at least one additional peripheral site are printed in black.

| Dataset | NP Likeness | Fsp$^3$ | FC$_{stereo}$ | nSPS | nPBF (*in silico*) | nPBF (protein-bound) |
|---|---|---|---|---|---|---|
| **Molecules exhibiting the highest descriptor value** | | | | | | |
| | F313 (+0.88) | F058/F312/F313 (1.00) | F058 (0.50) | F058 (50) | F312 (0.073) | F312 (0.061) |
| | F058 (+0.68) | F055 (0.92) | F055 (0.42) | F055 (44) | F313 (0.067) | F058 (0.060) |
| **Fragments** | F312 (+0.58) | F294 (0.77) | F312/F313 (0.33) | F312 (40) | F024 (0.060) | F313 (0.057) |
| | F322 (+0.56) | F184 (0.67) | F030/F032 (0.31) | F313 (34) | F189 (0.058) | F264 (0.055) |
| | F055 (+0.40) | F283 (0.62) | F225 (0.25) | F030 (33) | F005 (0.056) | F024 (0.052) |
| **Molecules exhibiting the lowest descriptor value** | | | | | | |
| | F264 (–1.82) | F009 (0.20) | F009/F283 (0.00) | F009 (10) | F009 (0.024) | F012 (0.019) |
| | F009 (–1.81) | F304 (0.23) | F102 (0.06) | F304 (12) | F138 (0.027) | F009 (0.022) |
| **Fragments** | F102 (–1.78) | F102/F012 (0.25) | F264 (0.07) | F186 (13) | F070 (0.028) | F304 (0.029) |
| | F001 (–1.39) | F001/F073 (0.33) | ..... (0.08) | F264 (18) | F073 (0.031) | F138 (0.030) |
| | F005 (–1.35) | F074 (0.36) | ..... (0.08) | F184 (18) | F102 (0.031) | F001 (0.032) |

| Dataset | NP Likeness | Fsp$^3$ | FC$_{stereo}$ | nSPS | nPBF (*in silico*) | nPBF (protein-bound) |
|---|---|---|---|---|---|---|
| **Molecules exhibiting the highest descriptor value** | | | | | | |
| **PDB** | Staurosporine [STU] (+1.06) | 8KB (1.00) | ATP/ADP/AMP/ANP (0.40) | 8KB (32) | 8KB (0.091) | 8KB/152 (0.083) |
| **PKIs** | Midostaurin (+0.66) | Gilteritinib (0.62) | Peficitinib (0.28) | Peficitinib (37) | Repotrectinib (0.054) | n.d. |
| **ChEMBL PKA** | CHEMBL3099612 (+1.84) | Sphingosine (0.89) | CHEMBL4080906 (0.27) | CHEMBL4080906 (44) | CHEMBL148333 (0.061) | n.d. |
| **ChEMBL Kinases** | CHEMBL1356390 (+3.93) | ..... (1.00) | CHEMBL23552 (1.00) | CHEMBL1596335 (61) | Propionaldehyde (0.150) | n.d. |
| **Oral Drugs** | Artenimol (+3.88) | ..... (1.00) | Kanamycin (0.83) | Memantine (64) | Cysteamine (0.132) | n.d. |
